# From biological data to oscillator models using SINDy

**DOI:** 10.1101/2023.08.25.554817

**Authors:** Bartosz Prokop, Lendert Gelens

## Abstract

Periodic changes in the concentration or activity of different molecules regulate vital cellular processes such as cell division and circadian rhythms. Developing mathematical models is essential to better understand the mechanisms underlying these oscillations. Recent data-driven methods like SINDy have fundamentally changed model identification, yet their application to experimental biological data remains limited. This study investigates SINDy’s constraints by directly applying it to biological oscillatory data. We identify insufficient resolution, noise, dimensionality, and limited prior knowledge as primary limitations. Using various generic oscillator models of different complexity and/or dimensionality, we systematically analyze these factors. We then propose a comprehensive guide for inferring models from biological data, addressing these challenges step by step. Our approach is validated using glycolytic oscillation data from yeast.

## Introduction

Many important physical, chemical, and biological systems are characterized by periodic changes, also called oscillations. Examples of such oscillatory systems are in physics the mechanical pendulum or alternating current, in chemistry the Belousov-Zhabotinsky (BZ) (1, 2) or Briggs-Rauscher (3) reactions, and in biology circadian rhythms (4), population dynamics (5, 6) or the early embryonic cell cycle (7).

Because of their ubiquity in a wide variety of systems, scientists have long been interested in understanding the underlying processes that describe and regulate the observed periodic behavior. To do this, they often use the language of mathematical models in the form of differential equations that can describe and predict the dynamics of a system. Such models are usually derived through experimental work, first principles, and/or scientific intuition (8).

Prominent examples are the derivation of the simple pendulum model from observation and first principles described by Huygens (9) and later by Newton (10), or the identification of the underlying mechanism of the BZ reaction by Field, Kórós and Noyes (2) and the formulation of the Oregonator model (11).

### Design principles of biological oscillators

As in biology, the biochemical regulatory network is typically not completely characterized, it is difficult to derive model equations from first principles. However, biological oscillators have also been found to follow a few key design principles (12). To date, all biological oscillators have three properties: **negative feedback, time delay**, and **nonlinearity**. Negative feedback is crucial to reset the system to its starting point after one oscillation period. Moreover, this feedback needs to be sufficiently time-delayed, and the interactions must be sufficiently non-linear, to avoid the system getting stuck in a stable steady state.

By combining these basic design principles with experimental measurements, many mathematical models have been constructed that are capable of describing biochemical oscillatory processes. However, there exist many different ways to incorporate the three key properties that are required to ensure oscillations. A time-delayed negative feedback can be implemented explicitly by including a time delay in the equations, leading to delay differential equations, which are mathematically quite difficult to study. Examples include the Mackey-Glass model for the variation of CO_2_ levels in blood (13) and the tumor suppressor protein p53 in response to DNA damage (14). Another way to include a time delay is to introduce many intermediate steps (thus increasing the dimensionality of the system), which is how time delays result in mass action type models (15, 16) or for simpler formulations as in the Fitzhugh-Nagumo (17, 18) and Goodwin models (19, 20) by the strength of nonlinear coupling of state variable equations in the system. Also, where the negative feedback is implemented can vary; it can e.g. act on the production of the oscillating variable, or on its degradation (21, 22). Furthermore, nonlinearity can be incorporated directly, either in the form of higher order polynomial terms (as in the Fitzhugh-Nagumo model (17, 18)), or nonlinear expressions such as ultrasensitivity in the form of Hill functions (as in the Goodwin model (19, 20)), or by increasing the amount of state variables (or dimensions) of a model (as in mass action type models (23, 24)).

This illustrates that formulating models using a combination of design principles, scientific intuition, and experimental measurements can potentially lead to ambiguous model formulations where the choice of specific model terms may be biased towards the scientist’s preference. Furthermore, in the era of big data in biology identifying underlying patterns and formulating such models ‘by hand’ may become increasingly difficult or impossible (25, 26).

### Data-driven model inference in biology

The recent emergence of data-driven methods, also called machine learning, has enabled the identification of such patterns in large data sets (27). Such methods can be divided into two main types (28): (i) *Black-box* methods that are trained on known data and are capable of predicting future outcomes, but do not provide interpretable models (model-free inference). (ii) *White-box* methods, which aim for interpretability and provide explicit mathematical models directly from data (model-based inference).

The focus of research in systems biology lies mostly in the identification of network structures (29), such as gene transcription networks or protein-protein interaction networks. Here one typically uses expression data, applying black-box methods based on neural networks (30) (e.g. Graph Neural Networks (31)) or spectral methods (29) (e.g. GOBI (32)). With these methods it is possible to understand the underlying structures of a biological system, e.g. causal interactions (33), but not the actual nature of interactions which lead to e.g. temporal oscillatory behavior.

To address the issue of interpretability while exploiting the ability of black-box approaches to identify patterns in measured data, mixed methods, also called gray-box approaches, have been developed. Examples of such methods include e.g. Physics-Informed Neural Networks in physics (34–36) or similarly termed Biology-Informed Neural Networks in biology (37), which aim to improve interpretability of results by intergrating prior knowledge into the design of trained neural networks. Other methods translate the structure of neural networks into interpretable mathematical equations via symbolic methods. For example, ‘Symbolic Deep Learning’ (38) aims at achieving this on the entire dynamical system, while ‘Universal Differential Equations’ (39) use neural networks only as a means to describe the most intricate, unknown parts of a dynamical system.

However, the simplest and most straightforward methods are white-box applications which are able to provide models of dynamical systems directly from data. Such methods are mainly regression-based and their prominent representatives are the ‘Nonlinear Autoregressive Moving Average Model with Exogenous Inputs’ (NARMAX) (40), ‘Symbolic Regression’ (41) and ‘Sparse Identification of Nonlinear Dynamics’ (SINDy) (42).

Especially SINDy has recently gained popularity in physics (43, 44), engineering (45), chemistry (46) and biology (47, 48) where it has shown great potential when applied on synthetic data. However, the SINDy method has been scarcely applied to real data. Examples include data from generic systems (48, 49), settlement data (50), gene expression data (51) and mainly experimental data from predator-prey dynamics (52– 54), where SINDy was able to identify relevant aspects of the respective biological systems.

### Structure of this work

Here we aim to determine why the original SINDy is not (yet) often used on (biological) experimental data (beyond predator-prey dynamics), with a particular focus on biological oscillatory systems. We start by ‘naïvely’ applying SINDy to three selected, experimental data sets of oscillatory systems. We do so considering limited prior knowledge, no noise filtering and using RIDGE regression similarly to the original publication by Brunton et al.(42)). We explore the simple pendulum, the chemical oscillations of the BZ reaction, and measurements of glycolytic oscillations in yeast. This allows us to define the main limiting aspects of the SINDy method. We then investigate these limitations using a set of commonly used, generic oscillator models varying in complexity and/or dimensionality: the Fitzhugh-Nagumo model, the Goodwin model, and a mass action type model of the cell cycle. Based on this study, we formulate specific mitigation approaches and propose a step-by-step guide for model identification of dynamic (oscillatory) systems in biology using SINDy or other regression-based data-driven methods. Finally, we apply our guide to the introduced example of glycolytic oscillations in yeast.

## Results

### Application of SINDy to experimental data

Before applying SINDy to experimental data, we briefly explain how we approach such model identification (see Fig. 1).

**Fig. 1.**
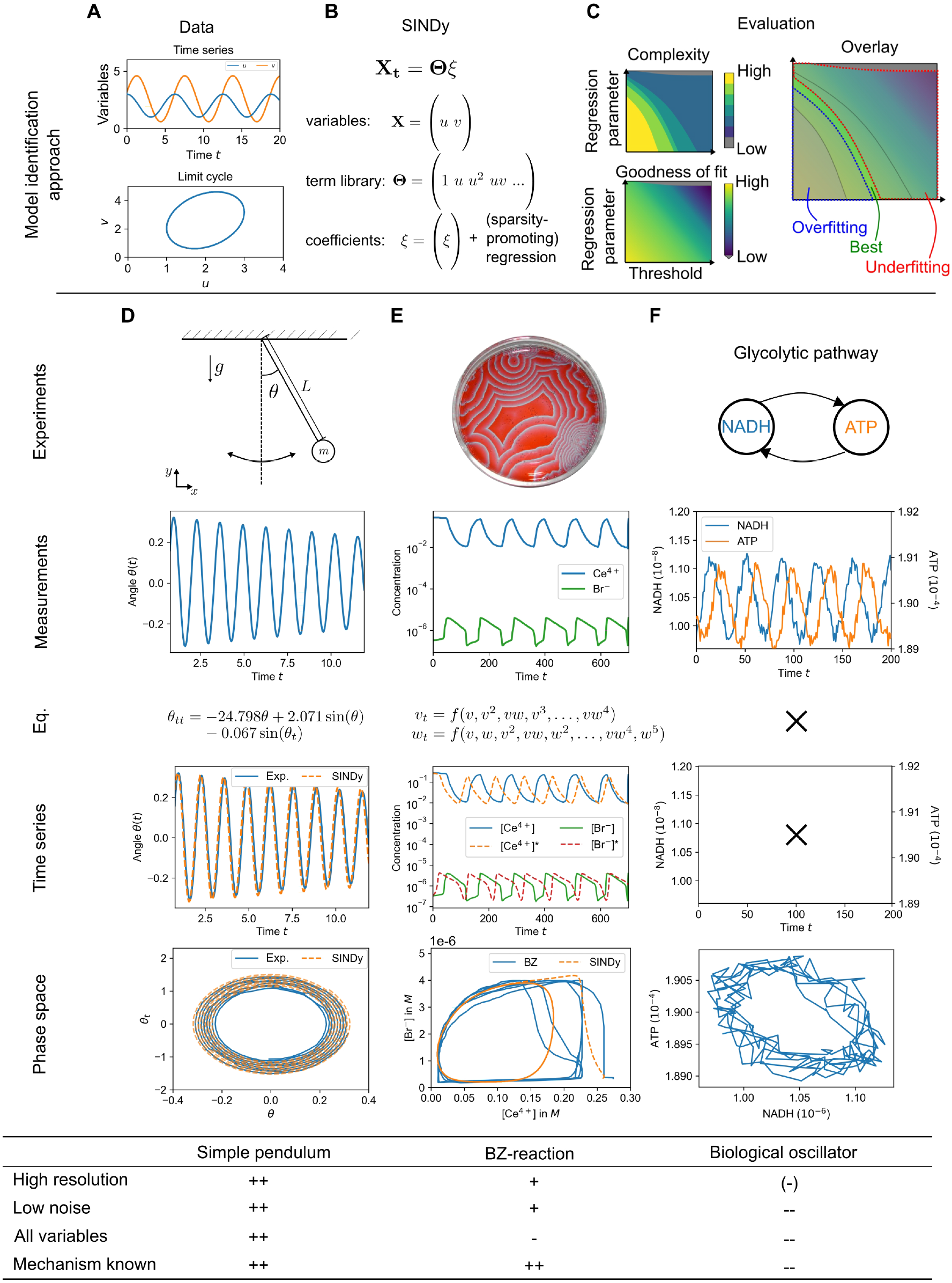
Model identification with SINDy from data of selected, real oscillatory system. **A**,**B** The SINDy method takes experimental data and creates a system of linear equations with a term library containing all possible terms and their coefficients. **C** Models are identified by applying a regression algorithm that solves the system. To determine the ‘best’ model, we scan through the hyperparameters of the regression algorithm and evaluate complexity (the amount of terms selected to be part of a suggested model by SINDy) and goodness of fit (here with the coefficient of determination or R^2^ score). By overlaying complexity and goodness of fit we can determine for which sets of hyperparameters the identified models, overfit, underfit or provide the best approximation to the data. In order to identify limitations of the SINDy method, we try to identify models of three selected oscillatory systems from experimental data: The simple pendulum, the Belousov-Zhabotinksy reaction, and oscillations in the glycolytic pathway. The systems and the acquired experimental data represent different levels of data quality, sufficient access to state variables, and knowledge of the underlying mechanisms. This can also be seen by applying the SINDy method to the data: **D** For the simple pendulum, we are able to determine the correct underlying system that reproduces the observed dynamical behavior. **E** For the BZ reaction, we are able to identify a model that captures the behavior but is not sparse, since it has to account for the nonlinear reduction of the system to only two variables. **F** Finally, when applied to data from glycolytic oscillations, we see that the SINDy method fails to identify a suitable model. (See Sup. notes 2 and 3)

We start by acquiring either new or existing data from the literature on an oscillatory system (see Fig. 1A). Using a single, measured time series of all available variables, we generate the target vector ***X*** (more precisely the time derivative ***X***_*t*_) of all variables and propose a suitable term library Θ containing all terms to be evaluated (see Fig. 1B, all libraries used in this manuscript can be found in Tab. 3 in the supplementary materials). This term library thus also contains the type of interactions between the different variables that we allow in the model, leading to a system of linear equations:

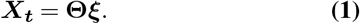

Finally, we apply a regression algorithm to this system, in our case RIDGE regression with sequential thresholding (55), and generate a family of models for different sets of hyperparameters (regression parameter and the underlying threshold). This approach enforces sparsity in models by cutting off all coefficients ***ξ*** and respective terms in the equation which fall below a preset threshold *l* or are excluded by varying the regression parameter *α*:

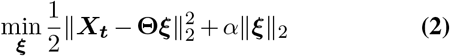

We classify the models by complexity (number of terms involved in the model), which we want to be as small as possible, and a goodness of fit measure, for which we use the coefficient of determination or *R*^2^ score (56) (see Fig. 1C). The *R*^2^ score quantifies how well a model is able to explain the observed behavior of the variables, where 1 is fully explanatory and 0 is not. In special cases, the *R*^2^ score can be negative, indicating that the model is not able to account for any dynamical behavior in the data.

We use both complexity and the *R*^2^ score at the same time, as high complexity correlates with high coefficients of determination or over fitting. However, we want to obtain the simplest possible model (low complexity) with the highest possible *R*^2^ score. We determine this by investigating the generated contour plots of complexity and *R*^2^ score (see Fig. 1C where we show over-fitted, under-fitted and the ‘best’ results from visual inspection) and determining the optimal result.

With this simple model identification approach, we set out to identify models from experimental data of oscillatory systems.

For this purpose, we choose three different systems with their respective data sets (see Fig. 1D-F): the simple pendulum, for which we acquired the data ourselves from video images; the well-known BZ reaction, for which we use existing data from the original publication by Field, Kórós and Noyes (FKN) (2); and time series of glycolytic oscillations between NADH and ATP acquired from Özalp et al.(57).

These three oscillatory systems not only represent three different fields (physics, chemistry, biology/biochemistry), but also differ in their quality in terms of resolution, noise, access to all state variables, and prior knowledge of the underlying mechanisms. The self-generated data of the simple pendulum represents one end of the spectrum, where we have high quality data and the mechanism is known. For the BZ reaction, the data quality is high, but not all state variables are measured. Finally, in the case of most biological oscillatory systems, such as glycolytic oscillations, we lack high quality data, only have information about some of the state variables, and do not have a complete picture of the underlying mechanisms that drive the oscillations.

These aspects are also reflected in the quality of model discovery using SINDy. For the self-generated, high-resolution and low-noise data of the simple pendulum (see Fig. 1D), combined with the knowledge of all variables (term library contains sin and cos, see Tab.3) and the underlying mechanism, the correct form of the ordinary differential equation is easily discovered:

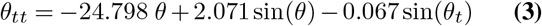

If we additionally apply the small angle approximation (i.e. *sin*(*θ*) *≈ θ*, if *θ ≪* 1) to reformulate the found equation into its well-known form, we see that SINDy provides us with the known equation of a damped simple pendulum. Additionally, we can also derive additional information of the underlying system, such as the length of the pendulum *L*, the friction parameter *b* or the mass *m* (for more details see Sup. Note 2):

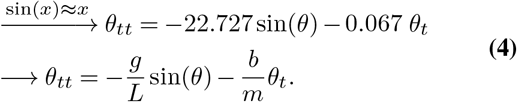

In the case of the BZ reaction, we have high quality (high resolution, low noise) data, but we only have access to two of the three relevant state variables (see Fig. 1E). The BZ reaction, first described by Belousov in 1959 (1), is a family of chemical systems that are an example of non-equilibrium thermodynamics and lead to the emergence of temporal or spatio-temporal oscillations. In their work, FKN studied the bromate-cerium-malonic acid system, where the main interacting compounds are the negatively charged bromine (bromine ions) [Br^*−*^ ], the positively charged cerium [Ce^4+^ ]*/* [Ce^3+^ ], and the neutral bromic acid [HBrO_2_] (2), which cannot be measured potentiometrically due to its neutrality. As a result, this chemical system that has been modeled with the three-component Oregonator model (58, 59) must be reduced to two components:

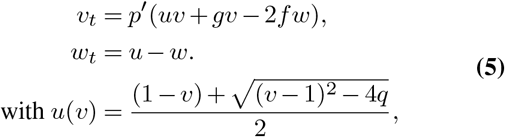

with *u* = [HBrO_2_], *v* = [Br^*−*^] and *w* = [Ce^4+^] */* [Ce^3+^] . Assuming that we do not have this knowledge about the underlying mechanism, applying SINDy with a term library containing only combinations of *v* and *w* will only be able to approximate the reduced system with higher order terms and thus models of high complexity. This results in a quantitatively correct but does not provide us with an interpretable model (see Eq. 7 in Sup. Note 3).

For the last case of glycolytic oscillations in yeast, we arrive at the other side of the spectrum, where provided data from Özalp et al.(57) is scarcely sampled and has high noise levels (see Fig. 1F, for more details see Sup. Note 8). Glycolysis has been intensely studied over the last 70 years (60, 61) and is a process that takes place in prokaryotic and eukaryotic cells in order to convert glucose to smaller substances. The main function is to provide the cell with adenosine triphosphate (ATP) when respiration does not take place (62). If dense suspensions of non-growing cells or cell-free extracts of the yeast *Saccharomyces cerevisiae* are starved, temporal oscillations can appear (61). The main driver behind glycolytic oscillations are dynamics of adenosine diphosphate (ADP) and ATP interactions (63), which form the basis of the most prominent model of glycolytic oscillations, the Selkov model (64):

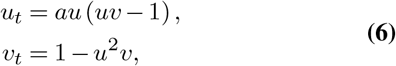

with the dimensionless *u* = [ADP] and *v* = [ATP].

Glycolytic oscillations are usually measured through the autofluorescence of the hydrogenated nicotinamide adenine dinucleotide (NADH) which oscillates in phase with ADP (65), as most chemical compounds taking part in glycolysis are optically silent (62). The data set we use, measures the oscillations of NADH, or indirectly of ADP and intracellular oscillations of ATP (57). However, SINDy is not able to identify suitable models in the form of the Selkov model (nor any other form) from the provided experimental data.

#### Challenges of model inference

From this naïve application of the SINDy method on experimental data, we identify four main challenges found in (biological) oscillatory systems:

### Insufficient resolution

Sufficient resolution plays a crucial role in model identification in general and for SINDy in particular; SINDy reportedly struggles in low-data scenarios (50, 66, 67), which has led to proposals to extend the method(49, 68, 69). However, the limitations of SINDy and the proposed extensions have not been sufficiently studied, and their proof of concept has been done on generic examples, such as the Lorenz system. Such systems usually do not reflect the dynamical behavior of real biological oscillators, which are often characterized by alternating slow and fast dynamics (12), resulting in pulse-like interactions of successive long/slow and short/fast phases called time scale separation, which can be difficult to capture if the resolution is insufficient. Alternative sampling approaches have been proposed here (70), but optimizing sampling strategies requires prior knowledge of the dynamics or may be impossible due to technical limitations.

### High noise

Besides low-data limits, SINDy has also shown reduced performance when the data are noisy (42, 66), since SINDy fits the derivative ***X***_*t*_ and not the actual measured time series, which can lead to non-unique solutions (71). Several ways of dealing with this have been proposed(49, 69, 72), and Delahunt and Kutz have developed a toolkit for noise handling when applying the SINDy algorithm (73). For the special case of biological transport models Lagergren, Nardini *et al*. (74, 75) have shown that using additional noise filtering via a neural network on (spatio)temporal data can significantly improve model identification with regression algorithms. However, the toolkit and other extensions also provide a proof of concept in dynamical systems that do not represent biological oscillatory behavior or experimental data, i.e., strong time-scale separation and/or insufficient temporal resolution. Thus, the limits of the methods in experimental situations are not fully known yet, and several extensions suggest the application of additional noise filtering (69, 73) without evaluating the impact of this choice on model identification.

### Number of dimensions

In order to derive correct models from data, SINDy needs information about all relevant state variables (also called dimensions) that are able to fully describe the dynamical systems. By ‘relevant state variables’, we refer to the smallest, finite number of variables needed for an unambiguous description of a dynamical system (76). Often such knowledge is not provided and either too many or not enough variables describing a system are measured, an example being the BZ reaction where one dimension is missing. Again, several extensions have been proposed that involve the use of auto-encoders (77, 78) or delay embedding (70, 79) to reconstruct dimensions or reduce the system. However, these approaches require high quality data to provide sufficient information to unfold missing dimensions or reduce existing ones, and therefore do not reflect data on dynamical systems in biology, where sometimes a variety of dimensions (e.g., fluorescent markers) are measured but suitable temporal data are lacking. Furthermore, in the case of high-dimensional systems we also encounter the so-called ‘curse of dimensionality’ (80), which obstructs the derivation of relevant information from such systems. Especially for biological oscillatory systems, this can pose significant challenges for parameter estimation, either with SINDy or other methods (81).

### Limited prior knowledge

Some of the above challenges can be overcome if there is prior knowledge about the underlying system. Such knowledge can be respecting basic physical laws, e.g. conservation of mass, or the implementation of already known interactions between certain actors of a system. When such information is available it is more likely that data-driven model identification will be able to identify physically-sound, interpretable models, but in many situations in biology, knowledge beyond basic physical laws is not available.

These challenges are always at least partially present in any experimental domain, but especially for experimental biological systems they can pose challenges for model identification. In systems such as the simple pendulum, which is a ‘simple’ example with high-resolution, low-noise experimental data and a substantial amount of prior knowledge, identification is easy: we know that we have to obtain the angle *θ* to identify the underlying model. However, for more realistic systems it becomes increasingly difficult (BZ reaction), if not impossible (glycolytic oscillations). Therefore, we aim to characterize the influence of low-data limits and noise when evaluating biological oscillatory systems with SINDy and to answer the following questions:

Q1. What are the general limitations for the identification of (biological) oscillator models with respect to data availability, resolution, and noise?
Q2. How does the choice of common noise filtering techniques affect the success of model inference for oscillatory systems?
Q3. What is the role of strong time-scale separation in biological processes in model recovery?
Q4. How do different nonlinearities in the formulation of biological oscillator models affect model inference?
Q5. How can model identification for high-dimensional oscillator systems work in low-data limits?

While answering the questions, we aim to address challenges faced by many experimentalist wanting to apply this method on their experimental data and provide a broad overview of relevant aspects that have to be taken into account when doing so.

### Impact of data availability, sampling, and noise on model inference from synthetic data

We begin our investigation by examining the availability of data (the amount of information provided in a time series), the sampling of data, and the influence and mitigation of noise. To address these questions and the other questions, we start with synthetic data generated from a set of generic, biologically motivated oscillator models that represent a non-exhaustive variety of model architectures in biology.

#### Generating synthetic data sets

Mathematical models used in biology can generally be classified by their dimension, or set of state variables, and the complexity of the interactions they describe, which in this case is the order of the terms included in the mathematical model (see Fig. 2). Thus, models in the model space of possible descriptions of a system can be seen as closer to biological reality if they use only simple lower order interactions obeying mass action kinetics and provide complex dynamical behavior through more dimensions. In contrast, models that take a higher-level approach and abstract high-dimensional systems with more complex interactions are a conceptual representation of biological reality. Both modeling principles have advantages: biological models allow us to directly relate certain behaviors to biological actors, but conceptual models allow us to more easily study the dynamical behavior of a system. Models that lie outside the model space are either overly complicated or overly simplified, and thus provide insufficient or redundant information.

**Fig. 2.**
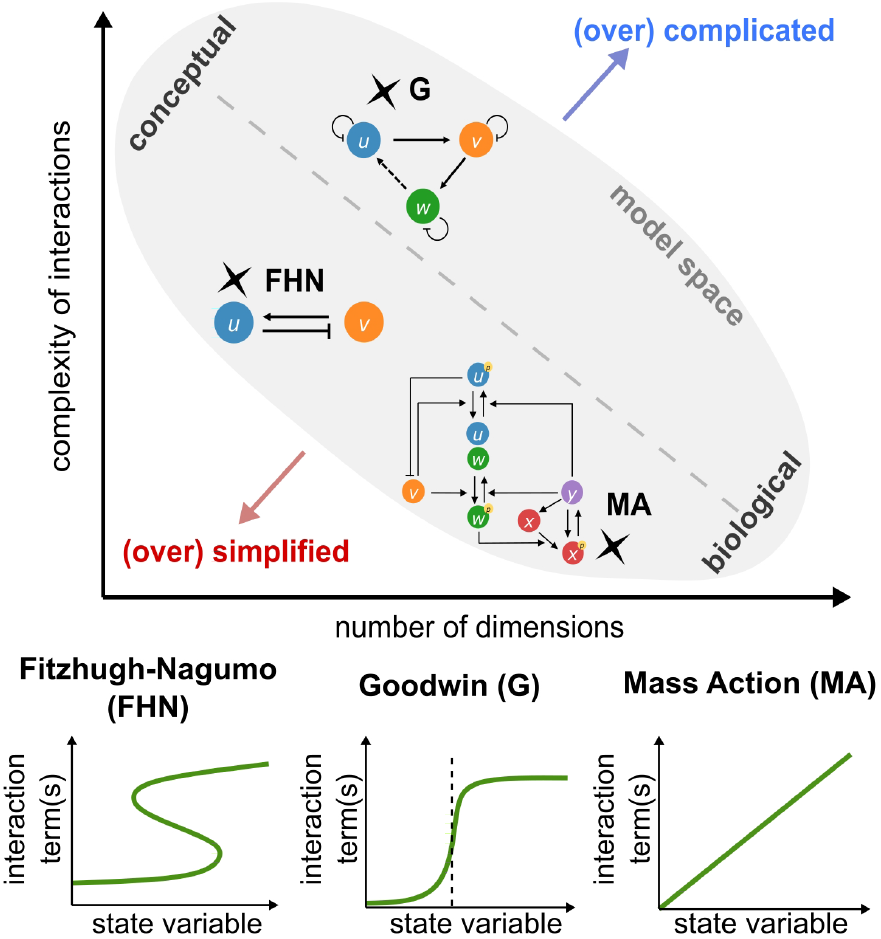
Classification of oscillator models in the model space of biological processes. Biological (oscillator) models can be characterized under two aspects: complexity of interactions and number of dimensions (state variables). There is a model space where both aspects are leading to functional models. If a model has low complexity of interactions and low amount of dimensions it is usually (over) simplified, or if both aspects are high it is (over) complicated. Models on the diagonal provide a trade-off between both aspects. In our work we consider three generic biological (oscillatory) models that can be applied to a range of biological systems: Fitzhugh-Nagumo oscillator (with cubic (bistable) interactions), Goodwin oscillator (ultra-senitivity in the form of a Hill function) and the (cell cycle) mass action model (with lower order interactions).

From this defined model space, we select three models that cover specific aspects and have been used to describe different biological systems:

### Fitzhugh-Nagumo

The Fitzhugh-Nagumo (FHN) oscillator was developed in the 1960s to describe the oscillatory spiking of neurons (17), but can also be used as a high-level description of other biological systems, such as cardiac dynamics (82), and early embryonic cell cycle oscillations (83). This model has a moderate complexity in its interactions between the variables, using a cubic nonlinearity instead of typical mass action dynamics. However, the FHN model allows oscillatory behavior in only two dimensions. We used the FHN equation of the following form:

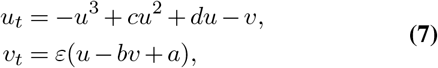

where *ε* is varied in the later stages (other coefficients are shown in Table 1). Here, the variables *u, v* correspond to a membrane potential and potassium and sodium channel reactivation (for neurons), excitation and recovery of cardiac cells (cardiac dynamics) or activity of the APC/C complex that inhibits the activity of Cyclin B-Cdk1 complex by degradation of Cyclin B, which is pushing a cell into mitosis (early embryonic cell cycle).

**Table 1.**
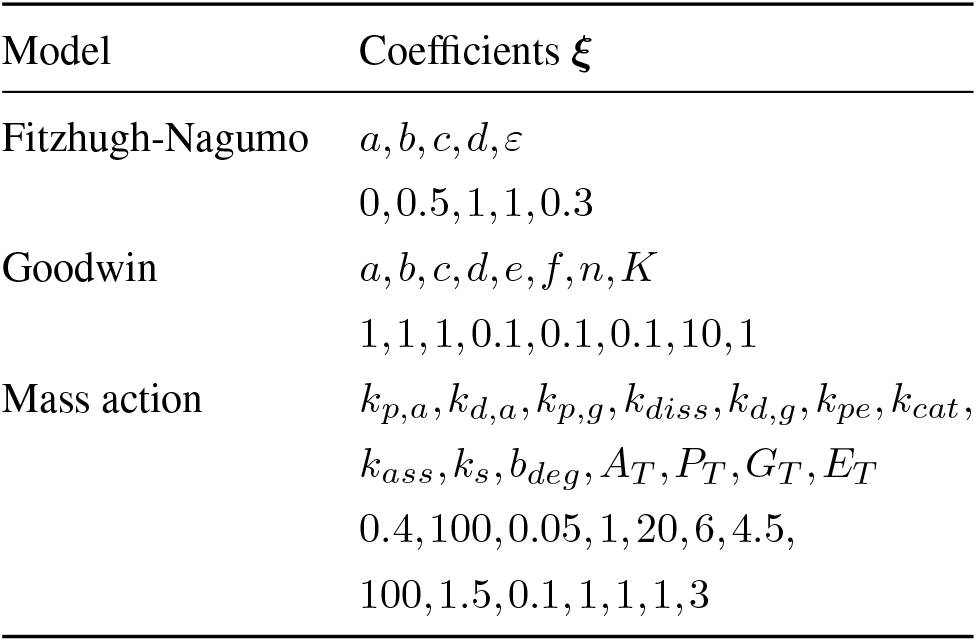
Coefficients of the respective model and the values used in this work.

### Goodwin

The Goodwin (G) oscillator was developed by B. Goodwin, also in the 1960s, in order to describe the phosphorylation/dephosphorylation processes of a transcription factor (19, 20) or circadian rhythms (20). This model has been widely extended and studied over decades and has a higher complexity of nonlinear interactions than the FHN oscillator through the nonlinear response curve in the form of a Hill function *H*(*x*), while avoiding any other higher order interactions between the variables. The delay of the negative feedback is implemented by an additional variable. Here we use the Goodwin oscillator in the following form,

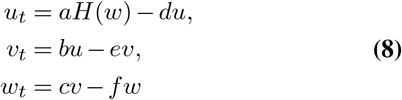

with the Hill-function defined in function of an arbitrary variable *x* as:

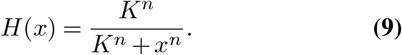

The coefficients can be found in Table 1. The variables *u, v, w* can be, e.g. interpreted as a clock-mRNA, clock-protein, and a repressor (circadian rhythm).

### Mass action

Mass action models are used to describe the dynamics of a wide variety of biological systems. They are characterized by more variables (higher dimensional), but simpler, mostly low-order interactions as their rate of change is directly proportional to the product of their activities or concentrations (84). Such interactions typically consist of linear or quadratic terms. In this work we use the cell cycle mass action model developed by Hopkins et al.(85), which describes the oscillations of the early embryonic cell cycle of *Xenopus laevis*, based on low-order (mass action type) interactions of six interacting molecules, avoiding complex descriptions of interactions in the form of ultrasensitive response curves or higher order terms (86). The model equations are as follows:

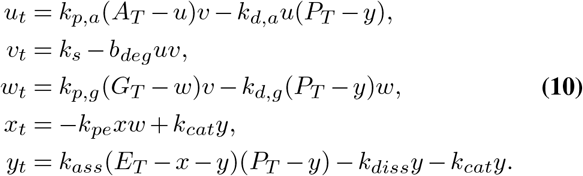

for which the parameters can be found in Table 1. Here, similarly to the FHN model for the cell cycle, the variables of *u* and *v* are interpreted as CyclinB-Cdk1 and APC/C activity.

In order to study model identification in experiment-like circumstances, we simulate one single time series of the respective models with a time step of *dt* = 10^*−*3^. We control the amount of information by changing the number of periods *T* included in the simulated time series *t*_sim_ = *n*_periods_*T* (see Fig. 3A). Using a fixed number of periods in the time series allows us to compare the results of different oscillatory systems despite their different period lengths. We further impose low-data limits by subsampling the simulated time series equidistantly to mimic typical experimental situations. To further resemble experimental situations, we add random (white) noise *η* which is normally ditsributed (Gaussian noise) to the simulated data sets consisting of fractions of up to 10% of the standard deviation *σ* of the signal itself^1^,

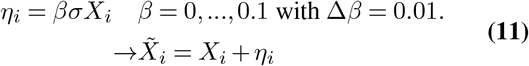

**Fig. 3.**
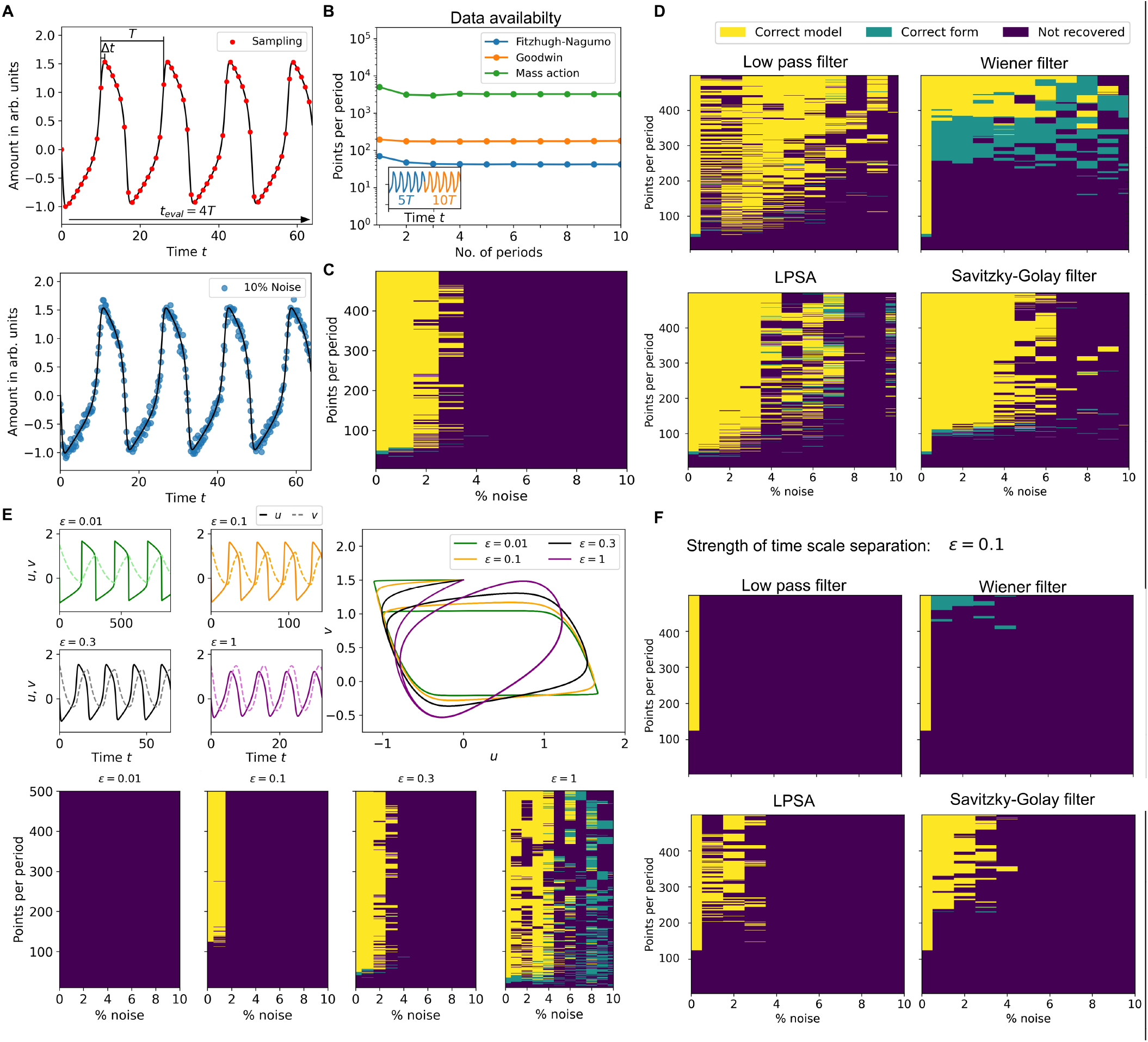
Investigation of data availability, equidistant sampling and added Gaussian noise for weak and strong time scale separation *ε* in the FHN model of Eq. 7. **A** The chosen models are simulated to generate one time series with *dt* = 1 *·* 10^*−*3^ which are later subsampled to simulate low-data limits. Noise is added in the form of up to 10% Gaussian noise. **B** By varying the length of the time series by the information content in information units (periods), we determine that increasing the length of the time series does not affect the amount of points needed for model recovery if at least 4 periods are present. **C** Combining simulated time series of length *t*_eval_ = 4*T* and added Gaussian noise, SINDy is able to recover the underlying model correctly (yellow) or its correct form (turquoise) for at least 45 points per period with up to 3% noise. **D** Noise reduction techniques (low-pass filter, Wiener filter, LPSA and Savitzky-Golay filter) are able to improve the performance of SINDy for higher noise levels, however the success depends on underlying methodology (more details in text). **E** An important aspects of biological oscillatory systems is time-scale separation, which can be changes with the *ε* parameter in the FHN model. For stronger time scale separation (small *ε*) identification requires high-resolution/low-noise data and fails for to strong separation. **F** The strength of separation also influences the performance of SINDy with noise filtering, where frequency filters (low-pass and Wiener filter) fail and techniques which estimate noise locally (LPSA and Savitzky-Golay) are superior.

It should be noted, that when scanning over the hyperparameters and using only a single time series can lead to varying model recover success (for a smaller threshold *l* identification is possible while for a slightly larger not) due to the variable nature of regression and the use of a random seed to generate the normally distributed noise. However, in Fig. S1 we show that despite this variability the overall outcome of our investigation is independent from the choice of seed.

#### Length of time series, equidistant sampling, and added noise

The evaluation of the length of the time series is important for the correct choice of experimental measurements, but also for the rest of the following study, since we choose the length of the time series to evaluate the performance of SINDy.

To do this, we simulate sets of time series for all models, varying the number of periods included: *n*_periods_ = 1 … 10. At the same time, we vary the number of points per oscillation period and determine the minimum sampling rate required to successfully identify the underlying dynamics (see Fig. 3B). We see that the required number of points per period (ppp) increases with the number of dimensions: The two-component FHN model requires at least 95 ppp, while the Goodwin and mass action models require 200 ppp (3 components) and 4500 ppp (5 components), respectively.

Furthermore, we see that the amount of ppp is reduced as more periods than one are included, but when the data set contains more then 2 periods increasing the number of periods involved does not influence the performance of SINDy. However, to avoid any potential influence on oscillations with strong time scale separation from the low number of periods, we decided to evaluate the performance of SINDy on time series of length *t*_eval_ = 4*T*.

In the following section we focus on data and noise bounds rather than dimensionality, we apply SINDy to synthetic data from the two-dimensional FHN model (Eq. 7). We define a successful identification, when SINDy selects all correct terms from the term library and identified coefficients lie within 2% of the true coefficients (marked in yellow, see Fig. 3C-F). If the coefficients lie beyond this threshold, SINDy was sometimes still able to recover the correct form of the underlying equation, thus providing important information on the system’s dynamics (marked in turquoise, see Fig. 3C-F). Otherwise, when model recovery was entirely unsuccessful, we colored this in dark blue (see Fig. 3C-F).

We start by applying SINDy to equidistantly sampled data from the FHN model, where we add different levels of Gaussian noise up to 10%. SINDy is able to correctly identify the underlying dynamics when there are at least 42 ppp and the added noise levels are low (below 2%, see Fig. 3C). This high sensitivity of SINDy to noise has already been reported in the literature (42, 49, 72, 73, 88–90). To handle such noise and improve performance, different noise filtering techniques were proposed (72), and more advanced methods were developed (49, 89, 90), which we will consider later.

#### Noise filtering techniques

Recently, in Cortiella et al.(91), an extensive analysis on the impact of selected noise filtering approaches for accurate model identification with SINDy has been conducted. Here we partially extend this analysis by specifically focusing on model recovery in biologically-motivated oscillatory systems. We select four different, commonly applied filtering techniques: A simple low-pass filter, a Wiener filter, a nonlinear filter based on local phase space averaging (LPSA) (92) and the Savitzky-Golay filter proposed in Lejarza et al.(72).

### Low-pass filter

The low-pass filter is a frequency filter that reduces the amplitude of high frequencies in a signal, which in the case of time series are the fast fluctuations of noise.

### Wiener filter

The Wiener filter is an enhanced low-pass filter, which assumes that noise is of high frequency and normal-distributed, which is the case for our added noise.

### LPSA

The local phase space averaging is a simple nonlinear filter, which averages the variability of measured variables in the phase space.

### Savitzky-Golay filter

The Savitzky-Golay filter regresses a low-order polynomial (e.g. third order) within a changing selected subset of data (moving window) in order to approximate the local changes while preserving the global trend.

The application of these methods generally leads to an improved model identification for the FHN model (see Fig. 3D). In particular, the low-pass filter was able to provide successful identification for high noise regimes because it is able to separate the normally distributed noise from the actual signal. However, for low-data and lower noise levels, the low-pass filter was not able to identify the model. This is a result of an imprecise cutoff frequency, which allows noise frequencies to be present in the time series. Selecting this cutoff frequency can be particularly difficult in low-data limits, as it can affect the shape of the oscillations and thus prevent correct model identification.

The Wiener filter consistently identifies the correct form of the underlying FHN model despite high noise levels when sufficient data is available (> 280 ppp). However, the recovery fails when less data is available, due to the Wiener filter’s fundamental assumption that the noise must be normally distributed: As the amount of data decreases, the identified error is no longer normally distributed, but uniform (we provide a detailed overview of the performance of noise filtering in Sup. Note 4, see Fig.S4).

The LPSA method is able to correctly identify the FHN model with the same number of points per period as the original SINDy and the low-pass filter. However, compared to the lowpass filter, it performs better in low-noise situations, while inconsistently identifying the dynamics at higher noise levels. As we show in Sup. Note 4, we see that this method is strongly dependent on the correct choice of delay in the approximation and is thus not able to filter stronger variability in high noise situations.

Finally, we studied the Savitzky-Golay filter, which was proposed by Lejarza et al.(72) to improve SINDy by applying a moving horizon (or window) optimization approach. We find that the Savitzky-Golay filter works well, but requires more data than low-pass filtering or LPSA, i.e. at least 100 ppp. For higher noise levels (above 6%,) using the Savitzky-Golay filter does not allow SINDy to correctly identify the underlying system. This is due to the low order polynomial approximation (regression) in moving windows: The higher the variability, the more precise must be the window size on which the regression is applied. Otherwise, the regression will overestimate the variability of the signal, resulting in a locally incorrect representation of the signal.

In conclusion, by choosing an appropriate (or wrong) filtering technique for the specific noise and data properties of the signal one can significantly improve (or hinder) model identification with SINDy. Even though all noise filtering techniques studied are capable of improving system identification, either too strong or too weak noise filtering can hamper data-driven discovery, as it can alter the shape of the oscillatory signal. This is especially important for biological oscillations that can contain multiple time scales, such as pulse-like behavior. Changing the shape of such pulses can then lead to a false description of the system.

### Different time scales in the system

Whenever a dynamical system is characterized by multiple time scales, this has general implications for regression-based model identification techniques such as SINDy. Time scale separation can be easily controlled in the FHN model by varying the parameter *ε*: for small *ε ≪* 1 the dynamics of *u* and *v* are strongly separated in time, resulting in relaxation (pulse-like) oscillations, while for large *ε ≈* 1 the dynamics are no longer separated and the oscillations become sinusoidal (see Fig. 3E).

In an equidistant sampling scenario, the strength of the time scale separation plays an important role for a successful model identification, as can be seen in Fig. 3E. Here we see our previously obtained results from Fig. 3B with the FHN and *ε* = 0.3. If we increase *ε* = 1, SINDy can identify the correct model equation for a smaller number of points (>15 ppp) and even in some cases for all tested noise levels (up to 10%). However, decreasing *ε* increases the amount of data required (*ε* = 0.3 – 42 ppp, *ε* = 0.1 – 116 ppp). and in the case of *ε* = 0.01 we are not able to recover the model with the provided amount of data (500 ppp). If the fast changes are not sufficiently sampled, the solutions will contain more spurious terms, despite the use of sparsity-promoting optimization algorithms.

In this context, we also examine how noise filtering can change the identification success in the presence of time scale separation. We only evaluate the success with the FHN model and *ε* = 0.1 (see Fig. 3F). Here, by applying a low-pass or Wiener filter, SINDy fails to identify a correct model in the presence of noise. This is again a consequence of the precise choice of cutoff frequencies, which can lead to the removal of high frequencies associated with the fast pulses, thus underestimating the strength of the time scale separation.

In this case, both the nonlinear LPSA method and the Savitzky-Golay filter are able to provide better results, although only for larger amounts of data (LPSA - about 200 ppp, Savitzky-Golay - about 230 ppp). The LPSA is able to handle lower levels of noise because it does not approximate the time series globally over time, but locally in phase space. With a sufficient number of points in time intervals of fast changes (horizontal and vertical edges of the limit cycle in phase space) and transitions to these phases (corners of the limit cycle in phase space, see Fig. 3E), the technique is able to capture the different time scales correctly. In the case of a Savitzky-Golay filter, the changes can be captured because again the fast dynamics of the oscillations are approximated locally within the moving window. Thus, with a sufficient temporal resolution, identification can be successful.

### Addressing noise and low-data in equidistant sampling with multiple trajectories

We see that identification of symbolic models of oscillatory systems is difficult when the data itself and its quality are modified to resemble experimental situations, even when we apply noise reduction techniques. Another suggested way to mitigate the problems of both low-data and high noise is to provide more than a single time series or trajectory.

Recently, two possible approaches have been proposed: The Ensemble-SINDy or E-SINDy extension (49), which applies bootstrapping, or more specifically bagging, to data and/or term library, thus synthetically increasing the amount of data that can be used to infer the underlying dynamics (for more details see STAR methods), and providing more trajectories, which has been shown to increase the success of model recovery (89, 90). Both approaches are conceptually similar in that they try to feed SINDy with more data from different situations, thus limiting the possible model space to models that can account for different trajectories.

We apply both approaches to data from the FHN model with *ε* = 0.3 in Fig. 4. It can be observed in Fig. 4A that E-SINDy performs worse than SINDy on the data provided, this in contrast to the results in Fasel et al.(49). For smaller amounts of noise, E-SINDy is not able to identify the underlying model, which in this case means that not all relevant terms appear in 90% of all submodels from the bootstrapped data (also called the inclusion probability *p*_inc_, which has to be larger than our chosen threshold probability *p*_thres_ = 0.9). We use this relatively high threshold compared to the original publication because in the case of unknown underlying model we are not able to choose an appropriate *p*_thres_ *a-priori*.

**Fig. 4.**
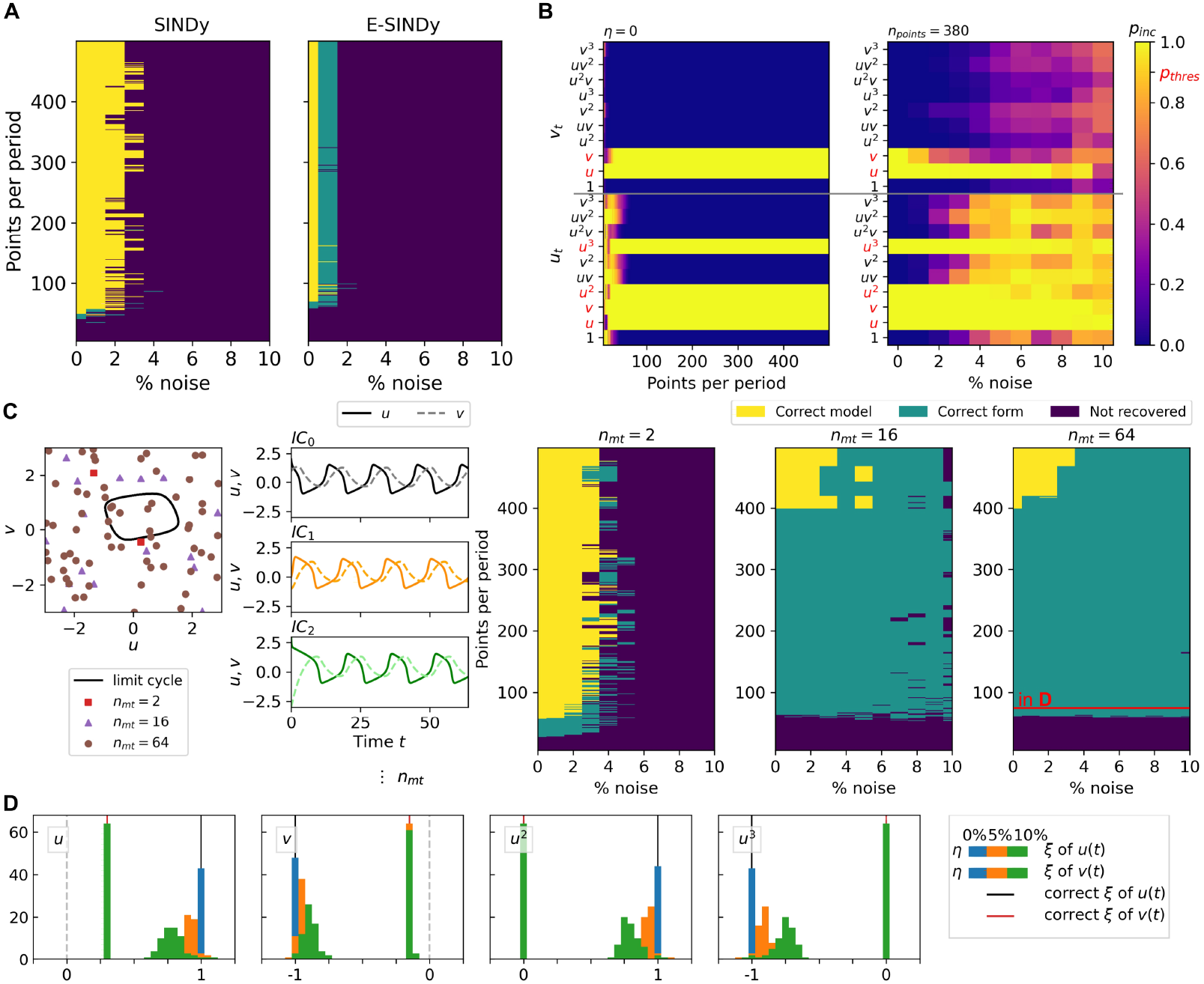
Addressing low-data and high noise with multiple trajectories. **A** Despite the claims of (49), the application of E-SINDy (right) compared to just SINDy (left) does not improve model recovery. **B** Inspection of the respective term probabilities to be included *p*_inc_ in the inferred model (threshold for inclusion in equation is *p*_thres_ = 0.9), show that E-SINDy underestimates the probability of several terms (e.g. *v* for *v*_*t*_) and overestimates mixed terms (e.g. *uv* for *u*_*t*_). **C** Providing more trajectories of the underlying dynamics with different, random initial conditions (IC) leads to an improved identification of the correct form of the FHN equation. The different IC are shown in the left panel: squares for two ICs, triangles for 16 ICs and circles for 64 ICs **D** Applying a subsequent nonlinear regression algorithm on noisy and sparse data with the identified models allows to identify the correct coefficients.

To understand the reduced performance, we examine the dependence of the inclusion probabilities on the amount of data and the noise level (see Fig. 4B). We see that without noise *η* = 0, E-SINDy needs at least 50 ppp to not only correctly select the terms, but also to reduce the amount of mostly mixed terms included in the inferred model of *u*_*t*_. Furthermore, the identification of the correct term breaks down when noise is included in the time series. Even at low noise levels, *p*_inc_ of the *v* term in *v*_*t*_ rapidly falls below the assigned *p*_thres_, while at high noise levels a wide range of non-relevant terms are included in the model. This begs the question why we are not able to achieve similar results as shown in Fasel et al.(49). The answer lies in the bagging method itself and the choice of data used in the original publication: Bagging (93), which stands for bootstrap aggregation, is a method that improves the predictive power of e.g. regression algorithms. However bagging is a smoothing operation that reduces the variance of the signal (94), which means that aspects such as time scale separation in limits of low-data and/or high-noise may be incorrectly smoothed and thus incorrectly approximated by SINDy. Indeed, in Fasel et al.(49), the signals either do not contain multiple time scales when studying a low-data limit, or large amounts of data are used if the variance is high. Thus, applying the E-SINDy method to the special case of biological oscillations may lead to unsatisfactory results due to bagging.

In contrast, increasing the number of trajectories available to SINDy without smoothing by bagging, as suggested by Ermolaev et al.(89), can allow to preserve the slow-fast changes in the oscillations. Indeed, we see that by introducing more trajectories of the same system with randomly chosen initial conditions (IC), we are able to significantly improve the performance of SINDy (see Fig. 4C). With two included time series, the minimum number of points is reduced to about 25 ppp, but the noise handling does not improve. If more trajectories are included (more than 8, Fig. 4C shows only a subset of the results), the required amount of data increases to 55 ppp, but SINDy is able to tolerate all introduced noise levels and can provide the correct form of the underlying model. The increased number of points is due to the variability introduced by multiple trajectories in low-data limits, which hinders the identification of the correct equation and simultaneously reduces the coefficient values when more data are available.

To address this issue, the models, identified with multiple trajectories, can be used as the basis for an adjacent simple parameter approximation where only the parameters of the identified model are optimized. This application leads to the correct identification of coefficients for low noise levels and reduced coefficients for high noise levels, thus improving the performance of SINDy for low noise levels (see Fig. 4D). With this we are able to improve the performance of SINDy without any additional noise reduction technique nor an increase in the sampling of the corresponding time series of experiments. However, such an approach does not show improved performance for stronger time scale separation than *ε* = 0.3. We have also obtained the results for the FHN with *ε* = 0.1, which can be found in Sup. Note 5, and despite the slightly improved handling of up to 6% additional noise, the number of points required is 2.5 to 8 times higher than for *ε* = 0.3 or when only SINDy is applied (see Fig. 3E).

### Mitigating time scale separation with improved sampling strategies

Another possibility suggested in Champion et al.(70) is the use of other sampling strategies, such as burst or specifically optimized sampling.

Therefore, we set out to evaluate whether optimizing sampling strategies is able to improve model identification in the presence of strong time scale separation. Here, we apply a different sampling approach that requires more knowledge about the underlying system than burst sampling, but should be able to provide more information about the dynamics: We keep the total number of points per period constant, but vary the distribution of points in the slow *δ*_*s*_ or fast *δ*_*f*_ changes of the time series, so that 1 = *δ*_*s*_ + *δ*_*f*_ = (*δ*_*s*1_ + *δ*_*s*2_) + (*δ*_*f*1_ + *δ*_*f*2_) (see Fig. 5A). The part of the time series assigned to fast changes is *α*_1_ + *α*_2_ = 0.2*T*, or 20% of the time period *T*, so if *δ*_*s*_ = 0.8, *δ*_*f*_ = 0.2 we get equidistant sampling. For a timescale separation of *ε* = 0.3, already increasing the number of points in fast changes improves the performance, with an optimal point at a 50*/*50 distribution (see Fig. 5B).

**Fig. 5.**
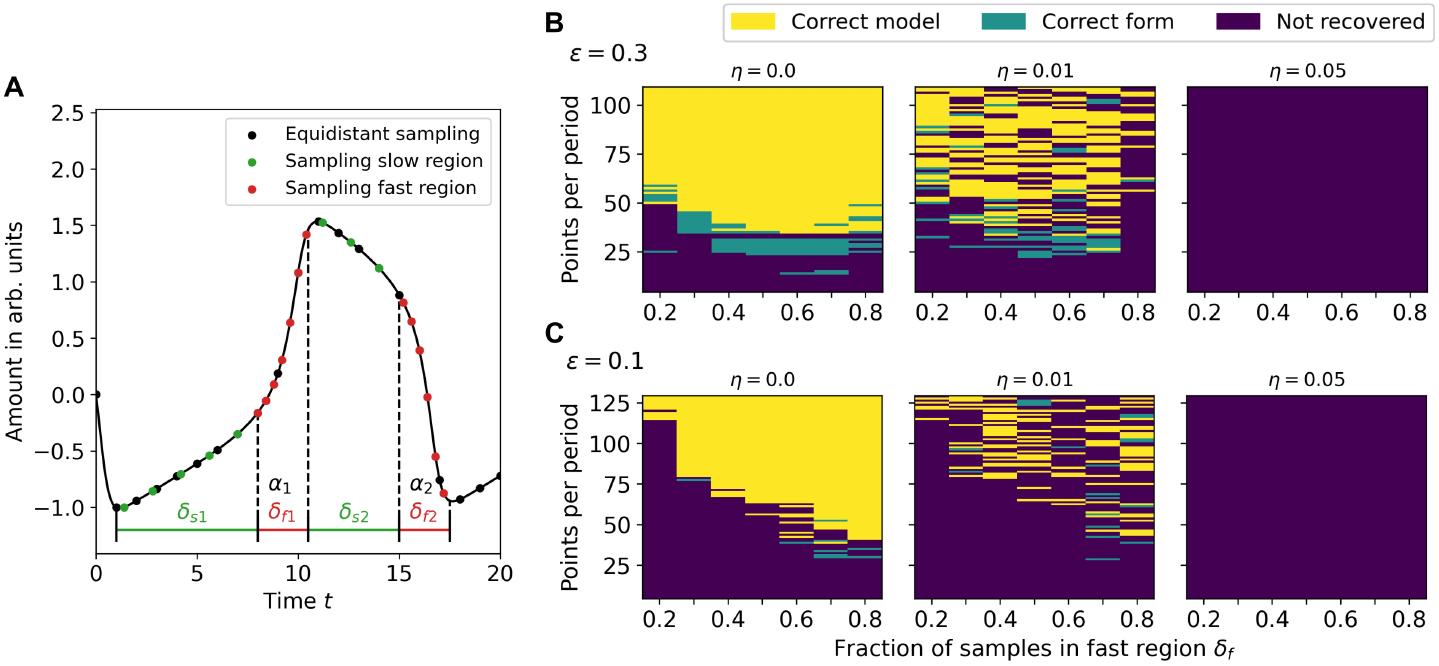
Mitigating time scale separation with optimized sampling. **A** The equidistant sampling can be optimized in order to provide more information on the fast dynamics by distributing the same amount of points unevenly between slow and fast changes. **B** For time scale separation of *ε* = 0.3 and by assigning 20% of the time series (*α*_1_ + *α*_2_ = 0.2*T* ) the performance is improved if already more then the basic 20% of points are evenly distributed in the fast changes, with an optimal distribution at 50/50 or 60/40 in favor of the fast changes. **C** For stronger time scale separation of *ε* = 0.1 an improved sampling strategy is able to significantly reduce the amount of points required, where the performance improves with the amount of points assigned to the fast changes, reducing the required amount from 115 ppp to 40 ppp with a distribution of 80/20.

The performance improvement is only successful for low noise levels (as shown in Fig. 5B), because computing derivatives in fast changes with small steps introduces more error due to increased variability, thus reducing performance.

However, for low noise levels, this method improves successful model recovery for even stronger time scale separation of *ε* = 0.1 (see Fig. 5C). By increasing the fraction of points *δ*_*f*_, we are able to reduce the minimum number of points required from 115 ppp with equidistant sampling to 40 ppp.

### Summary of the impact of data quality aspects on model identification (Answers to Q1, Q2 and Q3)

In the first part of the study, our goal was to answer what the sampling and noise limitations of SINDy are, and how do these limitations interact with the specific feature of time scale separation in biologically motivated oscillatory systems.

We find that successful recovery of the underlying models of oscillatory behavior depends more on the number of dimensions than on the number of periods provided in the time series. This has direct implications for experimental investigations, suggesting that high-frequency measurements should be preferred over long measurement times. Furthermore, we have shown that sufficient sampling is a key feature to recover underlying mechanisms and, as in most model identification, the more temporal data the better. However, more temporal data may not provide better performance when faced with high noise levels and/or strong time scale separation in the signal of oscillations (Q1). For the first part, we show that the introduction of noise filtering techniques can indeed provide a significant improvement in performance, but in low-data situations it can hinder identification if not set up precisely. In addition, when considering strong time-scale separation, noise filtering can significantly reduce the success of the investigation and can only provide improvement when large data sets are provided. Thus, even if data cleaning from experiments is able to provide a visually good signal, it may introduce artifacts that prevent identification of the underlying dynamics (Q2).

We have tried to avoid these aspects by studying the E-SINDy method and introducing more time series trajectories. Here we see that it is possible to avoid the application of noise reduction techniques and increasing sampling rates when there is a large number of trajectories. However, we show that the E-SINDy application, although undoubtedly applicable to low-data and high noise situations, is not suitable for relaxation oscillation data, since the applied bagging smooths the measured signal and acts as an unwanted signal filtering.

To enable identification for smaller time scale separations, we investigated whether an optimized sampling strategy could improve SINDy’s performance. Indeed, we were able to reduce the number of points to one third even for *ε* = 0.1 when equidistant sampling is applied. However, locally increased sampling, besides requiring prior knowledge about the system, only increases successful recovery when noise levels are low, since variability introduced in the derivatives on which SINDy regresses hinders correct identification of fast changes.

In general, the realistic case of strong time scale separation in experimental oscillatory data (high noise) is the most relevant limiting aspect when applying SINDy. Even if large amounts of data are available, identification can become either difficult or impossible (Q3), which makes SINDy (or the extensions studied here) applicable only to near-sinusoidal oscillatory systems, with important implications for its use in the field of systems biology.

### Influence of higher-order nonlinearities and high system dimensionality on model inference

After investigating the importance of data quality when using SINDy, we now turn to another important aspect of biological oscillatory systems: balancing the dimensionality of the model (number of components) and the complexity of the interactions between those components (see Fig. 2). While more complex higher-order nonlinear interactions can reduce the number of required state variables (lower model dimensionality), using many more components (higher model dimensionality) allows to use simpler low-order mass action interaction dynamics.

#### Ultrasensitivity in the Goodwin oscillator

In the FHN which we have used to study the data availability aspects, the non-linearities are introduced in the form of a cubic polynomial that allows us to be recovered using standard SINDy. However, some biologically-motivated oscillator models, e.g. the Goodwin model, chose to introduce nonlinearities in the form of ultrasensitive response curves that can be described by the Hill function shown in Eq. 9. For a three dimensional system (three interacting components) it has been shown that the required power of nonlinearity has to be at least 8: *n ≥* 8 (95). The standard SINDy method is not able to identify the Hill function or any other rational function from data, except if it can be approximated by higher order terms (as we could see in the case of the BZ reaction see Fig. 1 and Fig. S3 in Sup. Note 3) or if the functional form of the nonlinearity is provided. From Fig. 6A we see that if we generate time series with oscillatory behavior (*n* = 10), SINDy is able to approximate the behavior, but provides uninterpretable expansions of the Hill function (see Sup. Note 6) of the respective order. To address this problem multiple extensions have been provided during the last years, most notably the SINDy-PI (parallel implicit) which reformulates an equation with rational terms into an implicit form (96):

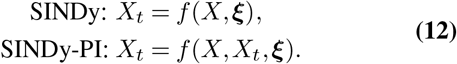

**Fig. 6.**
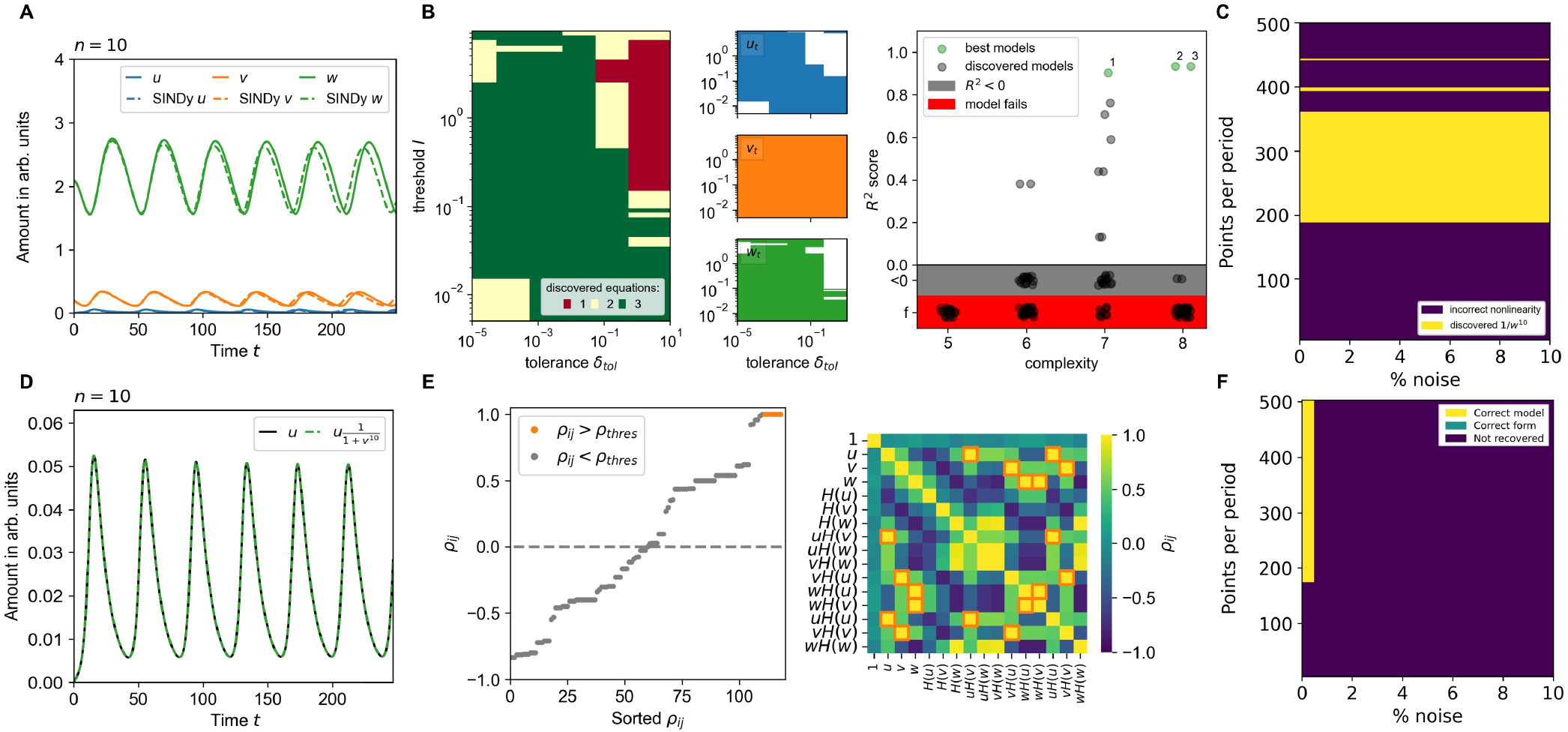
Enabling identification of a Hill-type response function with SINDy. **A** SINDy can not find the interpretable Hill-function but is able to approximate the nonlinearity (*n* = 10) with higher order polynomials. **B** SINDy-PI applies the SR3 regression (Eq. 14) using the hyperparameters threshold *l* and tolerance *δ*_tol_. Only for a subset of hyperparameters the full set of equations (*u*_*t*_, *v*_*t*_, *w*_*t*_) can be identified (see left panel). Using the *R*^2^ score a majority of models either does not reproduce the data or when simulated show non physical behavior (explosion of values). Three models with *R*^2^ *>* 0.9 (called best, shown in Tab. 2) provide the nonlinearity 1*/w*^10^ *≈ H*(*w*), but do not match the Goodwin model. **C** Identification of the Hill-function is highly sensitive to data amounts but not towards the amount of noise. **D** When the Hill functional form is known, SINDy is not able to identify the underlying model because of existent correlations between library terms, e.g. *u* and *uH*(*v*). **E** Correlations *ρ*_*ij*_ between different terms are identified and a threshold *ρ*_thres_ is applied from which correlated terms are excluded from the term library. **F** Excluding correlated terms allows SINDy to identify the correct underlying model. However, correlating the term library is only possible for low noise levels.

Thus the rational form of possible models is translated into a pure polynomial form and becomes solvable with the original SINDy approach. For the Goodwin model in Eq. 8, the first equation for *u*_*t*_ can be transformed into the following target for SINDy-PI (with *K* = 1 and a deliberately chosen *n*):

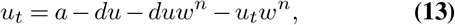

and by including the derivative *u*_*t*_ in the model identification process, the rational terms can be identified. However, the suggested SINDy-PI extension in Kaheman et al.(96) has only been shown to provide satisfying results when tested on synthetic data with more then 900 trajectories. As we have shown in the previous section, including more trajectories can indeed improve the identification notably. However, such large amounts of high-quality data are often not available in experimental setups, which has lead e.g. Brummer et al.(53) to reject the use of SINDy-PI in their work.

Nevertheless, the SINDy-PI extension provides the unique possibility to identify rational terms in a model within a sparsity promoting regression framework and we want investigate if SINDy-PI is able to discover either the underlying model or at least provide information on the rational nonlinearities in the Goodwin model. The original SINDy-PI uses STLSQ to promote sparsity, here we use the SR3 regression with *ℓ*_1_ regularization instead which is close to LASSO regression (97):

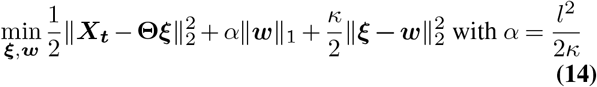

The SR3 regression introduces an additional auxiliary variable ***w*** which is used to relax the optimization problem. Because of the use of an *ℓ*_1_ regularization, the regression becomes sparsity promoting and is able to reduce coefficients *ξ*_*i*_ to 0. As a result the threshold *l* is no longer a strict threshold, but it is included in the choice of the regularization parameter *α*(*l*). Instead an additional hyperparameter, tolerance *δ*_*tol*_, is introduced. Although it is not directly included in the regression, it defines when the outcome of optimization is sufficiently good. Through the choice of hyperparameters, selecting optimal values for them is not intuitive compared to RIDGE regression.

Therefore we first determine which pair of hyperparameters provides all three equations (see Fig. 6B). Note that we use a single time trace (with 398 ppp) and a term library that only contains terms occurring in the Goodwin model (see Tab. 3). Using this set of models, we use the *R*^2^ score as a goodness-of-fit measure to identify which models provide the best fit. As can be seen from Fig. 6B, most models either are not able to describe the provided data (*R*^2^ *<* 0), or the model results in nonphysical dynamics (‘fail’ - *f* ) and only three models provide a *R*^2^ score higher then 0.9 (shown in Table 2). As can be seen, SINDy-PI is able to discover the true equations for *u*_*t*_ and *v*_*t*_ (with 1*/*(1 + *w*^10^) *≈* 1*/w*^10^), however it still struggles with the simpler equation for *w*_*t*_. Here SINDy-PI provides approximations of both equations which are able to qualitatively reproduce the dynamical behavior of the studied system (see Fig. S6). The correct discovery of the underlying model could be achieved by increasing the amount of data.

**Table 2.** Best discovered equations (*R*^2^ *>* 0.9 from Fig. 6B) with data from the Goodwin oscillator in Eq. 8 with no noise and 398 ppp using the SINDy-PI method with a library only containing terms already known from the model (compared with ground truth (GT).

|  | Equations |
| --- | --- |
| GT | $u_t = -0.1u + \frac{1.0}{1.0+w^{10}}$ $v_t = u - 0.1v$ $w_t = v - 0.1w$ |
| 1 | $u_t = -0.1u + \frac{0.03}{w^9} + \frac{1.0}{w^{10}}$ $v_t = u - 0.1v$ $w_t = -\frac{67.0v}{19.0w-100.0} + \frac{6.0w}{19.0w-100.0}$ |
| 2 | $u_t = -0.11u + \frac{0.03}{w^9} + \frac{1.0}{w^{10}}$ $v_t = u - 0.1v$ $w_t = \frac{50.0v}{17.0w+21.0} - \frac{5.0w}{17.0w+21.0} + \frac{0.5}{17.0w+21.0}$ |
| 3 | $u_t = -0.11u + \frac{0.03}{w^9} + \frac{1.0}{w^{10}}$ $v_t = u - 0.1v$ $w_t = \frac{50.0v}{17.0w+21.0} - \frac{5.0w}{17.0w+21.0} + \frac{0.5}{17.0w+21.0}$ |

However, even with a single time series, we are able to at least identify the correct type of nonlinearity (for high resolution, no noise data, small library). This knowledge can then be used to provide a better truncation of the term library for original SINDy. We therefore decided to identify how robust this identification is towards sampling and noise only focusing on how well one can recover the nonlinear Hill function (or its approximation 1*/w*^10^), see Fig. 6C. Here, SINDy-PI shows to be highly dependent on sampling, where sufficient resolution is required in order to identify the correct nonlinearity, but when more data is provided identification fails. This results from more ambiguous model identification of the nonlinearity when actual sparsity promoting regression methods are used, which we found not to be dependent on noise (see Fig. 6C). At this point, it is important to truncate the SINDy-PI library correctly, as the use of larger libraries can reduce successful identification of the nonlinearity (see Sup. Note 6).

More recently, SINDy has also been used to identify a model describing cancer and CAR T-cell dynamics(53), or population dynamics within an aphid-ladybeetle system(52) using extensive prior knowledge on the nonlinear interactions to provide these in the term library. Beside using SINDy-PI or extensive prior knowledge, another suitable way to determine the nonlinear rational term, is to provide new knowledge about possible nonlinear interactions between components of a biochemical system, which is usually gained from perturbation experiments. See for example the approach for obtaining a model of the tumor suppressor protein p53 in response to DNA damage in Geva-Zatorsky et al.(14), and of glycolytic oscillations in yeast cells in Nielsen et al.(65).

Assuming that we have the knowledge of the nonlinearity in Eq. 9, the Hill function can be introduced as an additional term in the term library in SINDy. However, using SINDy in this way does not lead to the correct identification of the Goodwin model. The reason for this is the random correlation of several terms contained in the library, e.g. *u* and *uH*(*v*) or *uH*(*u*) (see Fig. 6D). If these terms are present in the optimization problem, the regression algorithm is able to freely choose any coefficient for these terms, since they are almost identical and thus indistinguishable for the problem, e.g. for *u* and *uH*(*v*):

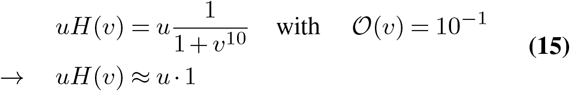

As a result, after initializing the library, it is important to identify any existing correlations between library terms. Two library terms are correlated if the result of the Pearson’s product-moment coefficient is *ρ* = 1. However, there is no common threshold *ρ*_thres_ above which two variables are correlated. We therefore decided to find a *ρ*_thres_ empirically by observing the distance of the term correlations *ρ*_*ij*_ from 1 (see Fig. 6E). By applying correlation, we are able to identify the culprits in the library and achieve successful model recovery when a sufficient amount of data (195 ppp) is available (see Fig. 6F). However, it should be noted that this method is highly sensitive to noise and requires precise noise filtering, which in itself is a challenge, as we have shown previously.

#### High dimensionality in the mass action oscillator

For the two models studied so far, we only considered two or three dimensions, where nonlinearities are explicitly expressed either by cubic terms or by the Hill function. In the case of the five-component mass action model the increase in dimensions is able to account for nonlinearity and delay. However, as we have seen from our investigation of experimental systems, despite a simpler representation and a closer relationship to an actual biological system, such an increase can effectively interfere with data-driven model identification.

Applying SINDy to the full dataset of a single time series, we observe that more data are required to correctly identify all the underlying equations compared to the previously studied models (2150 ppp in Figs. 3B and 7A).

Furthermore, we find that the identification with all dimensions present in the data is sensitive to noise and is not successful for our model even when large amounts of ppp are present (see Fig. 7A). Increasing the dimensionality of a system may decrease the ability to uncover the underlying dynamics. This is especially true when we have multiple variables acting on scales that are several orders of magnitude apart. As we have pointed out before, these limitations could be potentially overcome if the data set would contain more then one trajectory; however, providing tens or hundreds of trajectories of adequate quality for a high-dimensional system can be challenging.

**Fig. 7.**
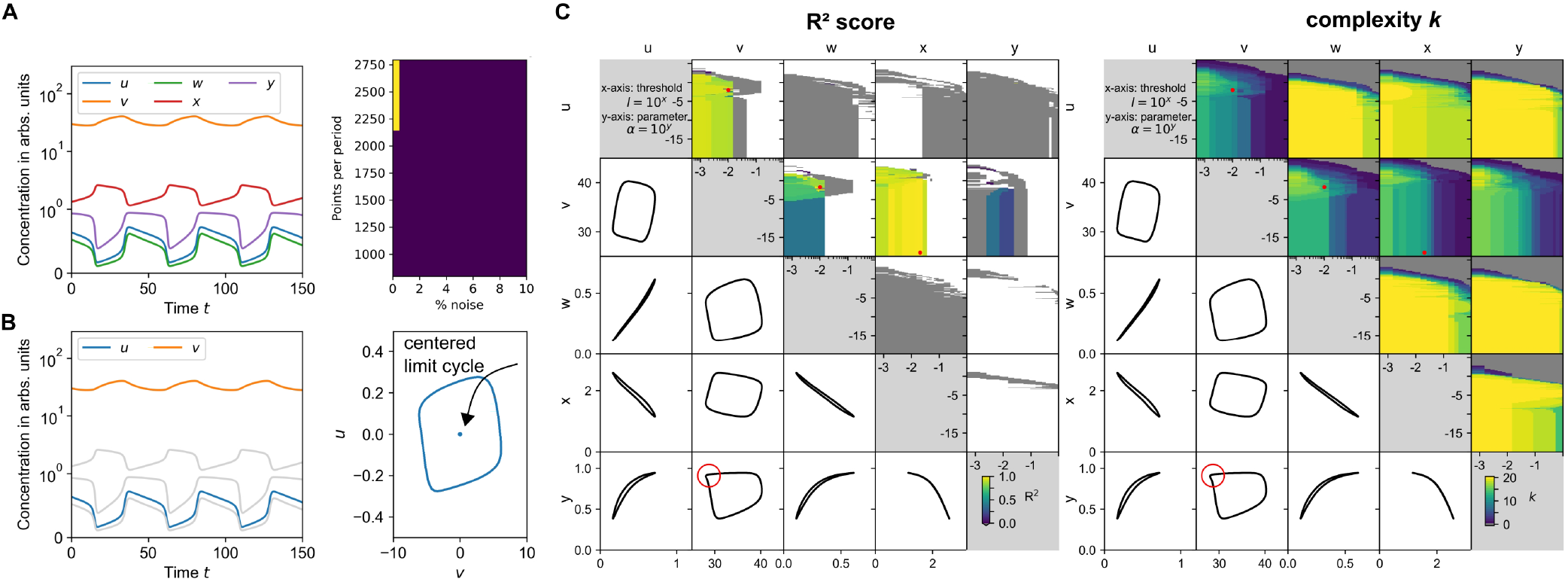
Identification of high-dimensional system by reducing set of state variables. **A** Model identification with high-dimensional data of the mass action oscillator requires high-data/low-noise situation in order to correctly identify the underlying system. **B** Reducing the system to only two variables, e.g. *u* and *v*, and centering the data around the point (0,0), assuming that the unstable fixed point is symmetric towards the limit cycle can be used to derive important dynamical information while reducing data requirements. **C** Applying SINDy to different combinations of state variables *u, v, w, x, y* and determining the R^2^ score (left) and complexity *k* (right) within a threshold *l* and regularization parameter *α* scan provides us with a set of models with high R^2^ scores and low complexity (red dots) that can be further studied dynamically and experimentally. These models occur when the state variables creating the limit cycle are well-separated thus providing sufficient information about the dynamics of the system. Only if the behavior can not be approximated by a cubic polynomial as for (*v, y*) (red circle) SINDy struggles to identify a suitable model, similarly as in the BZ reaction.

Thus, identifying such high-dimensional systems from all possible states seems highly impractical, even if all relevant variables are measured. We could reduce the number of state variables to the most relevant ones, e.g. by using encoders(77, 78). However, as shown especially in Chen et al.(78), large amounts of high quality data are needed to identify the most relevant aspects of an observed dynamical system, while the set of variables we provide is also the relevant set of state variables for the model.

At this point, we have shown that if there is not too much time-scale separation in the data, it is more advantageous to work with lower dimensional data, since a lower sampling resolution is required and SINDy is less sensitive to noise. This would move us away from a more biological model towards a conceptual, higher level representation of the dynamical system (as shown in Fig. 2). Even though we would not uncover the actual interactions, we could learn and understand the underlying mechanisms of the system.

When studying the BZ reaction (see Fig. S3 in Sup. Note 3) we have seen that not every reduction of a system is able to provide an interpretable model. Since we have access to all variables of the system, we can test the model identification of every pairwise combination of the state variables *u, v, w, x, y*. To do this, we center the limit cycles of the various reduced systems around (0,0), since for most generic models the unstable fixed point lies at (0,0) within the limit cycle, and we try to reduce the amount of additional terms that account for an offset (see Fig. 7B). We then scan the hyperparameters threshold *l* and the regularization parameter *α* over several orders of magnitude and determine the complexity *k* and the R^2^ score of the identified models in Fig. 7C. For the scan, we see that we are able to obtain high R^2^ scores for three combinations: (*u, v*), (*v, w*), and (*v, x*).

In this analysis, we also plot the projected phase space plots of the respective combinations, and we clearly see that when the variables forming the limit cycle are well separated, SINDy is able to identify important dynamical features of the system from the phase space. The only exception is the combination of (*v, y*) for which SINDy is not able to identify models leading to a high R^2^, despite a clear separation in the variables. This is a situation similar to the identification of the BZ reaction in the previous section. Since we have a sharp change in two interacting variables, this change cannot be approximated by a cubic polynomial. For the the mass action oscillator, such sharp changes are shown with a red circle in the phase plot of (*v, y*) in Fig. 7C, and similar behavior is found for the BZ reaction in Fig. 1E. In the case of the BZ reaction, polynomials of order 7 or higher were required to obtain a sufficient, but uninterpretable, approximation of the system, and for the mass action oscillator we do not provide such high order terms. This indicates an unfavorable reduction of the mass action system in the (*v, y*)-space and it requires either an additional variable or knowledge of the type of reduction that leads to this behavior.

However, when comparing the results of the R^2^ score with the complexity *k*, we choose the values of the threshold *l* and the regularization parameter that give the best trade-off between complexity and accuracy for the other well-separated variable combinations.

For (*u,v*) Eq. 10 is able to reproduce the dynamical behavior of the reduced system (see Fig. S8A, both in Sup. Note 7). Furthermore, the equations are remarkably close to the FHN equation in Eq. 7 and can even be analyzed analytically. With this reduction we are able to create a conceptual understanding of the underlying mass action system without the need for high quality data or the evaluation of all state variables at once.

Similarly the Eqs. 11 and 12 for the (*v, w*) and (*v, x*) respectively, are able to approximate the corresponding behavior in the reduced system (see Figs. S8B and C, both figures in Sup. Note 7). Both equations do not resemble a simple FHN-type equation, since all equations contain terms up to cubic order. Furthermore, as recently pointed out(98), a system consisting of two coupled bistable switches, i.e. the nullclines of a two-dimensional system are described by cubic polynomials, may lead to more robust oscillations and may be closer to actual biological reality.

#### High-order nonlinearities and high dimensionality in model identification (Answers to Q4 and Q5)

In the second part of the study, we tried to answer whether different formulations of nonlinearities in oscillatory models beyond the polynomial can be identified using SINDy, in this case evaluating the Hill function.

Since SINDy by itself is not able to provide rational terms, i.e. the correct form of the Hill function, it can only provide approximations of the nonlinearity in polynomial form, which may not be interpretable without prior knowledge. Therefore, we evaluated the extension SINDy-PI while applying it only on a single time trace, which reformulates the optimization problem with rational terms in an implicit form. When applying SINDy-PI with a library that is truncated to only contain terms included in the Goodwin model with the order of the nonlinear Hill-function *n* = 10 and on high resolution data (398 ppp) with no noise, we are able to discover a set of models that are able to reproduce the provided data. These models are also able to identify the approximated Hill-function (as 1*/w*^10^, but they struggle to provide the correct form for the simpler third equation *w*_*t*_. Nevertheless, SINDy-PI can be used to determine (if data quality and library truncation are suitable) possible nonlinearities within a model (here *H*(*w*) in *u*_*t*_). This knowledge can be then used to introduce suitable terms into the library for the original SINDy method to uncover the full correct model. Another possibility to discover nonlinearities using SINDy requires additional measurements, i.e. perturbation experiments, or extensive prior knowledge to determine nonlinear interactions and provide them within the term library.

We tried this by providing the correct Hill function formulation in the library, which led to an unsuccessful identification, from which we deduced that it is necessary to perform a prior correlation analysis between the library terms. This allowed us to identify possible correlations and thus enable successful model identification. Thus, we have shown and emphasized the importance of truncating the term library by correlation for regression-based model identification (Q4).

We then investigated the representation of time delay and nonlinearities in a higher-dimensional system with low-order interactions by applying SINDy to synthetic data from the five-component mass action model. As shown, the identification of this system requires high quality data, since SINDy is not able to handle additional noise in addition to a large number of points per period.

As a result, in the previous part we suggested that especially low-data limits and certain noise levels can be overcome by using lower dimensional systems. Using the existing set of relevant state variables, we attempted to infer models of a reduced two-dimensional system by cross-testing all variable combinations.

We find that identification is successful when the state variables are well separated in phase space and SINDy is able to unambiguously identify dynamical features.

It can be concluded that the identification of high-dimensional systems with SINDy, even with simple low-order interactions and access to all relevant variables, is challenging and requires high quality data. To enable identification, it is advisable to study possible reduced systems from which, depending on the separation and possibly available prior knowledge (to provide good approximations of terms in the reduced system), one is able to learn and even interpret analytically the dynamical behavior using a high-level representation (Q5).

### A step-by-step guide to applying SINDy on experimental data

After studying various limiting aspects of SINDy, we now summarize our findings in the form of a step-by-step guide when aiming at model identification with SINDy from oscillatory experimental data in biology. With this guide, shown in Fig. 8, we aim not only to systematize our approach to model identification, but also to provide a guide for experimentalists who want to better understand their dynamic biological system through models, without requiring expertise in classical model identification.

**Fig. 8.**
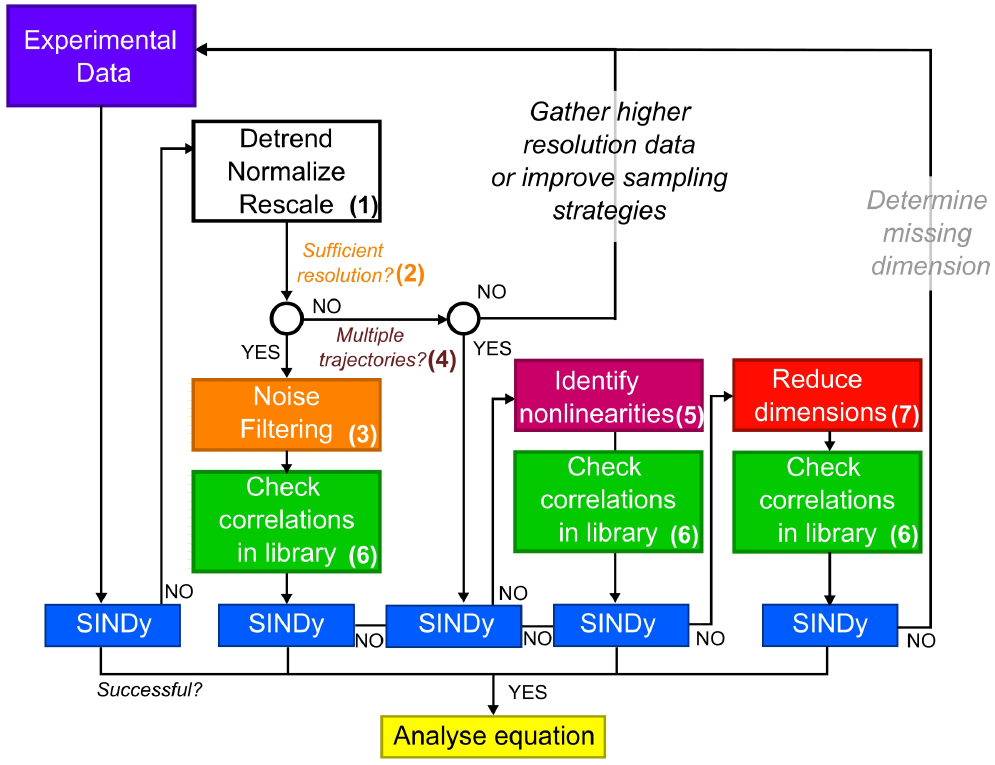
Overview of suggested step-by-step approach for application of SINDy on biological oscillator data. Resulting from the study on synthetic data, we suggest a step-by-step guide on how to approach model identification with SINDy when handling oscillatory experimental data in biology.

The steps included in the setup start with a first application of SINDy on experimental data, which is followed by (numbered as in Fig. 8):

1. classical data manipulation techniques not discussed in this work, namely detrending, normalization and rescaling of time series data (92).
2. investigating if provided experimental data meets minimal requirements towards temporal resolution (see Section ‘Length of time series, equidistant sampling, and added noise’ and following).
3. correctly filtering noise without influencing the outcome of model identification (see Section ‘Noise filtering techniques’).
4. suitable inclusion of more experimental data sets, either by measuring additional time series or optimizing sampling strategies (see Sections ‘Addressing noise and low-data in equidistant sampling with multiple trajectories’ and ‘Mitigating time scale separation with improved sampling strategies’).
5. investigating if nonlinearities can be identified using SINDy-PI for only one time series, and then implementing it in the original SINDy approach (see Section ‘Ultrasensitivity in the Goodwin oscillator’).
6. appropriate handling of correlations within the term library when an ultrasensitive response from the system equations is included in the term library (see Section ‘Ultrasensitivity in the Goodwin oscillator’).
7. an investigation of possibilities to reduce a high-dimensional system of state variables (see Section ‘High dimensionality in the mass action oscillator’).

If identification is unsuccessful despite following all suggested steps, the reason likely lies in the lack of knowledge about important missing state variables or interactions in the underlying dynamical system: Missing state variables or (nonlinear) interactions can make the identification of interpretable models difficult or even impossible if they cannot be well approximated within a reduced state space. In this situation, other techniques to uncover missing dimensions of a system have to be applied.

Nevertheless, using this approach we now return to one of the first experimental examples, namely glycolytic oscillations in yeast, for which we could not identify an underlying mathematical model (38 ppp, *dt* = 1*s*, see Fig. 1)^2^. We start by shifting the center of the attractor to (0,0), similar to the evaluation of the mass action model, and rescale the data with a min-max normalization (step (1), see Fig. S9B in Sup. Note 8 and Fig. 9A). These steps improve the identification, but do not lead to an interpretable model. Thus, we turn to one aspect of our study and apply a noise filtering technique, in this case a low-pass filter, since the oscillations do not show strong time-scale separation and are mostly sinusoidal (steps (3), see Figs. S9C and D). With this application, we are able to identify the following model at *α* = 1 *·* 10^*−*17^ and *l* = 1 *·* 10^*−*2^ with R^2^ = 0.7824 and complexity *k* = 15:

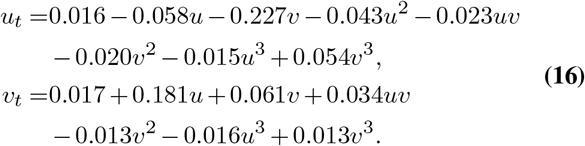

**Fig. 9.**
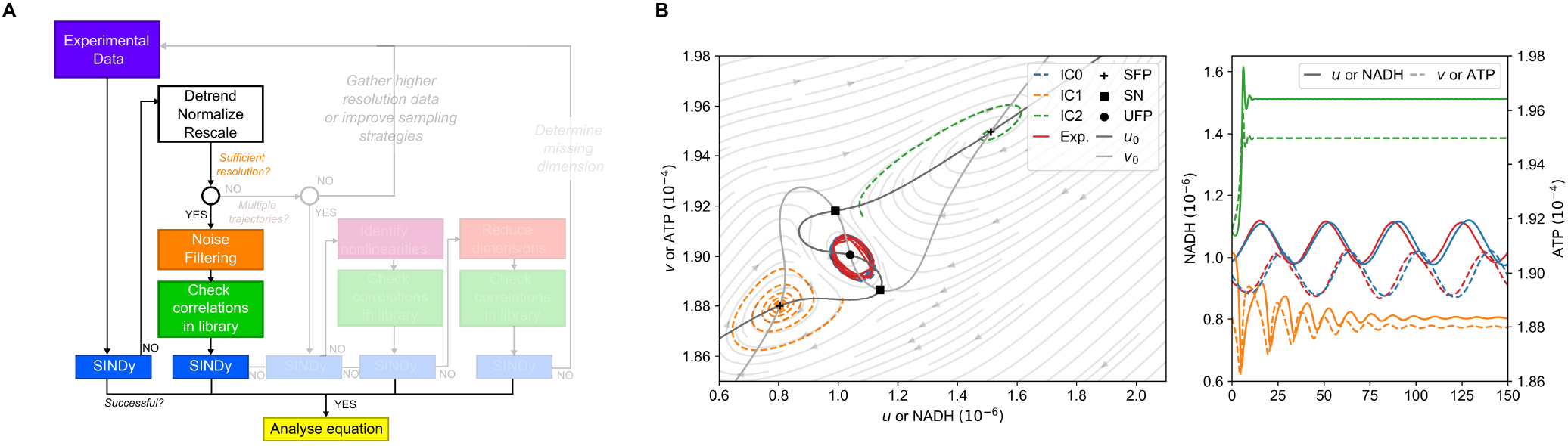
Enabling model identification from experimental data on glycolytic oscillations in yeast. **A** To enable model identification for glycolytic oscillations in yeast, the data are detrended, normalized and centered around (0,0) in phase space. These steps allow us to identify a model which is able to describe the dynamics (Transformed data not shown here). **B** The found model in Eq. 16 with a R^2^ = 0.7824 and a complexity of *k* = 14 is identified, which is able to describe the dynamical behavior and uncovers steady states in the system that are found in experiments but not represented in the popular Selkov model (Data has been transformed back to be compared to original experimental data; data in right plot uses the same color scheme as the left plot).

Remarkably, the identified model contains two equations up to the third order, which in turn can be compared to a model consisting of bistable interconnected switches as proposed by Parra-Rivas et al.(98). Due to its form, the equation is not easy to interpret or to study analytically. However, we study the equation numerically and are able to retrieve the null-clines and visualize the behavior in phase space (see Fig. 9B). Compared to the well-studied two-component model of glycolytic oscillations, the Selkov model in Eq. 6, we see that we identify a more complex description of the interactions, as the equation generates five fixed points (see Fig. 9B): An unstable fixed point (UFP) around which the limit cycle of oscillations occurs; two saddle points (or saddle nodes - SN) for high NADH and low ATP, and low NADH and high ATP concentrations/activities; two steady states (stable fixed points - SFP) for low NADH and low ATP, and high NADH and high ATP concentrations/activities. With this, we identify a biologically/biochemically sound model that is able to address the limitations of the Selkov model, already pointed out by Selkov himself (64): He points out that his simple model is unable to account for experimentally identified steady states, although these have already been described for oscillations in the yeast *Saccharomyces carlsbergensis* (99). Our data-driven model accounts for this behavior even though we provide almost no prior knowledge.

However, our model shows high complexity making it difficult to be interpreted analytically, which as a result from our study we assume to be due to the low temporal resolution (only 38 ppp, step (2)). Therefore, following our decision tree, we must either collect more additional data or improve the resolution of the experimental data used for model identification (step (5)). We suspect that increasing the resolution (either experimentally or even synthetically) could lead to an even simpler model (steps (4) and (5)).

#### Enhancing identification by synthetically increasing the resolution - an outlook

Since we don’t have access to more data, we wanted to try out what impact the application of a simple linear interpolation to our data and synthetically increasing the number of points would have on model identification (step (5)). Increasing the number of points by simple interpolation can be done in this particular case because the oscillations do not have strong time scale separation, however we do not evaluate the influence of interpolation on data-driven identification. We increase the number of points by a factor of 10, together with noise filtering, centering and normalizing of the data (see Fig. 10 and related to it Figs. S9E (increasing resolution), F (noise filtering), G (normalizing), H (normalizing and noise filtering)), and we identify a set of equations that preserves the same dynamical behavior as in the low resolution case while reducing the complexity from *k* = 15 for Eq. 16 to *k* = 12 in Eq. 13 (for more details see Sup. Note 8).

**Fig. 10.**
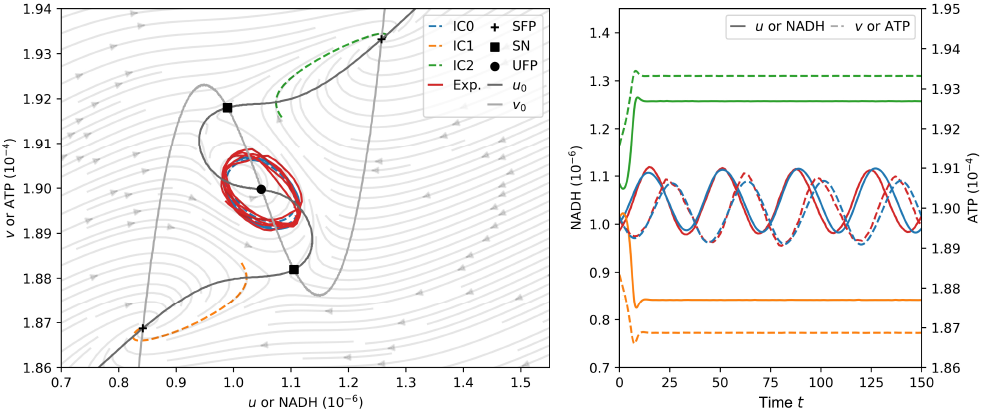
Improving model identification with interpolation of experimental data. Increasing the resolution by 100 times of the original amount of ppp (38 ppp to 3800 ppp) the identified model preserves the dynamical behavior while becoming not only easily interpretable but can also be studied analytically (Data in right plot uses the same color scheme as the left plot).

As a results, we assume that by increasing the resolution by a factor of 100 (ca. 3800 ppp, *dt*_2_ = 0.01, step (5)), we are able to achieve an even better approximation. As can be seen, we are able to identify a set of different models that represent the highest R^2^ scores and thus reproduce the experimental data increasingly well. Here we choose an equation that gives the best trade-off between R^2^ score and identified complexity *k*, which we find with hyperparameters *α* = 1 *·* 10^*−*17^ and *l* = 2 · 10^−2^ :

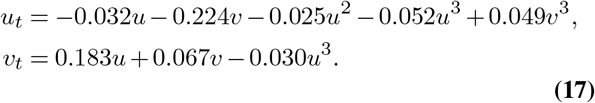

Evaluating how generalizable and/or closely related the model actually is to biological behavior is beyond the scope of this work. However, this analysis shows that by overcoming low-data limits, in this case by using interpolation, the results can lead to accurate and dynamically interesting and qualitatively relevant models that are interpretable. This aspect is an interesting line of work that was shown by Goyal et al.(69) to improve the performance of SINDy when a Runge-Kutta scheme is applied to the input data. However, to our knowledge, there has not been a comprehensive study of this topic in the context of SINDy.

## Discussion

In this work, we evaluated why, despite its recent popularity, the SINDy approach for data-driven model identification has rarely been applied to experimental data and has led to the identification of new models directly from experimental data. We focused on the identification of models describing oscillatory systems in biology, which are challenging systems for SINDy due to several factors. We have identified the main challenges by applying SINDy to self-generated or existing experimental data, which are insufficient resolution (or more generally, data availability and quality), high noise levels in experimental measurements, the number of state variables (or dimensions) that fully describe a dynamical system, and limited prior knowledge.

We investigated how the first three aspects in particular can influence the outcome of model identification in the special case of biological oscillatory systems. We selected three different oscillatory models (Fitzhugh-Nagumo, Goodwin, and a mass action model) capable of representing these challenges and investigated the performance of SINDy.

We have also shown that SINDy identification depends on the correct choice of noise filtering technique, and if chosen incorrectly, can obstruct correct identification. To mitigate this, we evaluated the performance of generating more data by either bootstrapping or including more experiments in the analysis and showed that for the special case of oscillatory systems, boostrapping can reduce SINDy performance and including more data is not able to mitigate strong time scale separation well. At this point, we have also investigated the choice of optimized sampling strategies and have shown that this can improve identification to some extent, although again it does not allow identification for strong time scale separation. We conclude that SINDy struggles especially when the data resolution is low and the oscillatory behavior is characterized by strong time scale separation. Here, identification requires very high resolution or may even be impossible, which is a crucial limitation of SINDy when applied to biological oscillatory data.

We then considered the specific aspects of nonlinearity and time delay in biological oscillatory systems, which are represented in the model either by high-order and complex expressions or by an increase in the number of dimensions (state variables). Here we have shown that for the Hill function commonly used in biochemical models, that the specially developed SINDy extensions SINDy-PI is able to provide knowledge on the approximated nonlinearities when data quality and library truncation is sufficient and SINDy only approximates the behavior with a high-order polynomial. Therefore, when using SINDy, prior knowledge about such high-order interactions must be provided either from SINDy-PI or from additional experiments and the resulting term library must be evaluated for possible correlations between terms to ensure correct identification. For high-dimensional systems, we have shown that even when no noise is applied, SINDy requires large amounts of data to correctly determine the identification. Thus, we have proposed and shown that cross-testing all available data sets of the state variables can enable the identification of a high-level representation of the dynamics in a low-dimensional system. As can be seen from our previous analysis, SINDy needs less data and can handle noise better to provide a suitable model. With this, we were able to identify a set of models that could even be dynamically studied.

All these findings have been condensed into a step-by-step guide for model identification with SINDy that can be applied to biological experimental data and beyond. Using our elaborate guide, we were able to enable the identification of an improved model for glycolytic oscillations directly from the original data set. By pushing the resolution by simple interpolation, we identified an analytically interpretable model of interconnected bistable switches directly from the data. This highlights the need for high resolution data, which is the bottleneck of model identification in general, but especially in biology. Therefore, model identification with SINDy has not yet led to the discovery of new interpretable models. In many cases, increasing the resolution cannot be achieved experimentally, and as we have shown, interpolation can play an important role.

In conclusion, we have identified the main limiting aspects of model identification with SINDy when applied to (biological) oscillatory experimental data and propose a step-by-step guide that can improve the identification success. Thus, with this work we aim to encourage the application of SINDy by biological experimentalists and increase the success of model identification directly from experimental data. However, different aspects require further studies to improve the performance of SINDy on experimental data, such as correct normalization and rescaling of data, the generation of more data without additional experiments through interpolation, and the discovery of hidden variables from experimental data.

## Limitations of study

In our study we apply simple intepolation in experimental data of the gylcolytic oscillator. However, data interpolation is a challenging task in data-driven model identification and SINDy, as if applied incorrectly it can alter the data to represent the result of the interpolation and not the underlying dynamics.

Another important aspect that we have not touched on is the correct positioning of the attractor or limit cycle in phase space. In our work, for the sake of simplicity, we centered the limit cycle around the point (0,0), either by removing the offset from the data or by normalization, which in many generic oscillatory models (e.g. FHN) is the unstable fixed point around which oscillations arise. However, even for the FHN model, the point is not positioned symmetrically within the limit cycle, and as we show in other work (100), correctly positioning the limit cycle in phase space is important for successful model identification.

Furthermore, we did not investigate the identification of hidden variables or dimensions directly from experimental data, e.g. in the case of the BZ reaction. The identification of such variables according to the Takens theorem (101) and delay embedding can significantly improve the success of data-driven model identification(70, 79) and result in successful model identification from experimental data(53).

Beside this, we also did not delve into other possibilities to tackle the challenging task of identifying strong nonlinearities like the Hill-function in a dynamical system. An example of such methods are ‘Universal Differential Equations’ (39) that can be combined with SINDy, to learn ‘simpler’ (low-order) interactions directly from data and use the strength of neural networks to discover the structure of strong nonlinearities such as the Hill-function.

Lastly, we did not carry out a detailed study of SINDy extensions that avoid using derivatives, such as Weak SINDy (43, 102). Here, the authors apply the weak numerical formulation of the problem, which can improve model identification for noisy or time scale separated time series data. We confirmed this as well, see Fig. S10. Therefore, a further study of non-derivative based SINDy or other methods that have shown better performance would be useful. These methods could improve model identification for oscillatory systems compared to derivative based SINDy methods.

## STAR methods

### Key resource table

#### Resource availability

##### Lead contact

Further information and requests for data or code should be directed to and will be fulfilled by the lead contact, Bartosz Prokop.

##### Materials availability

This study did not generate new unique reagents.

##### Data and code availability

- All experimental or synthetic data have been deposited at RDR by KU Leuven(103) and GITLAB (104), and are publicly available as of the date of publication. DOIs are listed in the key resources table.
- All original code has been deposited at RDR by KU Leuven (105) and GITLAB (104), and is publicly available as of the date of publication. DOIs are listed in the key resources table.
- Any additional information required to reanalyze the data reported in this paper is available from the lead contact upon request.

### Experimental model and subject details

This study did not include experiments with a specific model or subject.

### Method details

#### The SINDy method and its derivatives

In this work we use the original *Sparse Identification of Nonlinear Dynamics* (SINDy), developed by (42), and extensions of it, which are implemented in the PYTHON package PYSINDY (106, 107). SINDy is based on the assumption that dynamic systems can be described through differential equations in the following form:

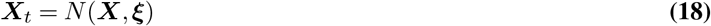

with a variable ***X*** = (*u, v*, …). The time derivative ***X***_*t*_ is a function of the variables ***X*** itself, combinations with other state variables and a set of parameters. Differential equations of this form can be also linearly combined, e.g. for component *u*:

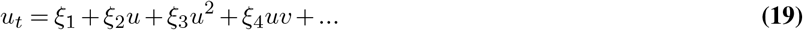

This equation can be rewritten as a row vector containing all combinations and derivatives of the quantity, called the term library and a parameter vector ***ξ*** containing all parameters:

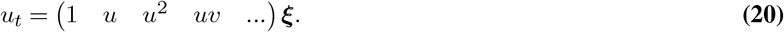

The values of each term in the library can be calculated from a single shot at a given point in time. If this system is extended to all available time points, a linear system of equations with the unknown parameter vector ***ξ*** and the term library matrix Θ is formed:

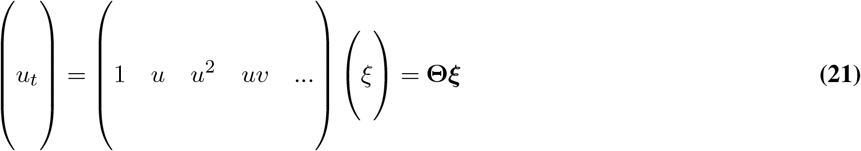

This system poses an over-determined optimization problem for values of ***ξ*** and can be solved by using sparsity promoting regression algorithms. In our work we use the RIDGE regression with sequential thresholding optimization algorithm (see Eq. 2). We evaluate the performance of SINDy and its extensions mostly using one trajectory (except for the section ‘Addressing noise and low-data in equidistant sampling with multiple trajectories’) with a set random seed to ensure that our results are reproducible.

#### The E-SINDy extension

Aside of this we also include the recently developed ensemble SINDy extension (E-SINDy) from (49). Here, the original system of linear equations in Eq. 21 is divided in to sets of subsystems, also called bootstrapping. The system can be sub-sampled in two different ways, either row/data wise only including subsets of measured data or column/library wise only including certain terms into sub-libraries into the optimization problem.

After solving all the subsystems, the solutions are aggregated and terms or their respective coefficients are assigned with a probability of inclusion. The authors of this approach claim that through bootstrapping application the model identification is more robust in low-data and high-noise regimes and therefore poses an optimal extension for our analysis.

For the different experimental systems under investigation, suitable term libraries are defined which can be seen in Table 3. The choice of libraries is motivated by prior knowledge (trigonometric functions for the simple pendulum, higher order terms in BZ reaction) or the expected behavior (3rd order terms for the glycolytic oscillations in assumption of a FHN type equation and from the Selkov model (64)).

**Table 3.**
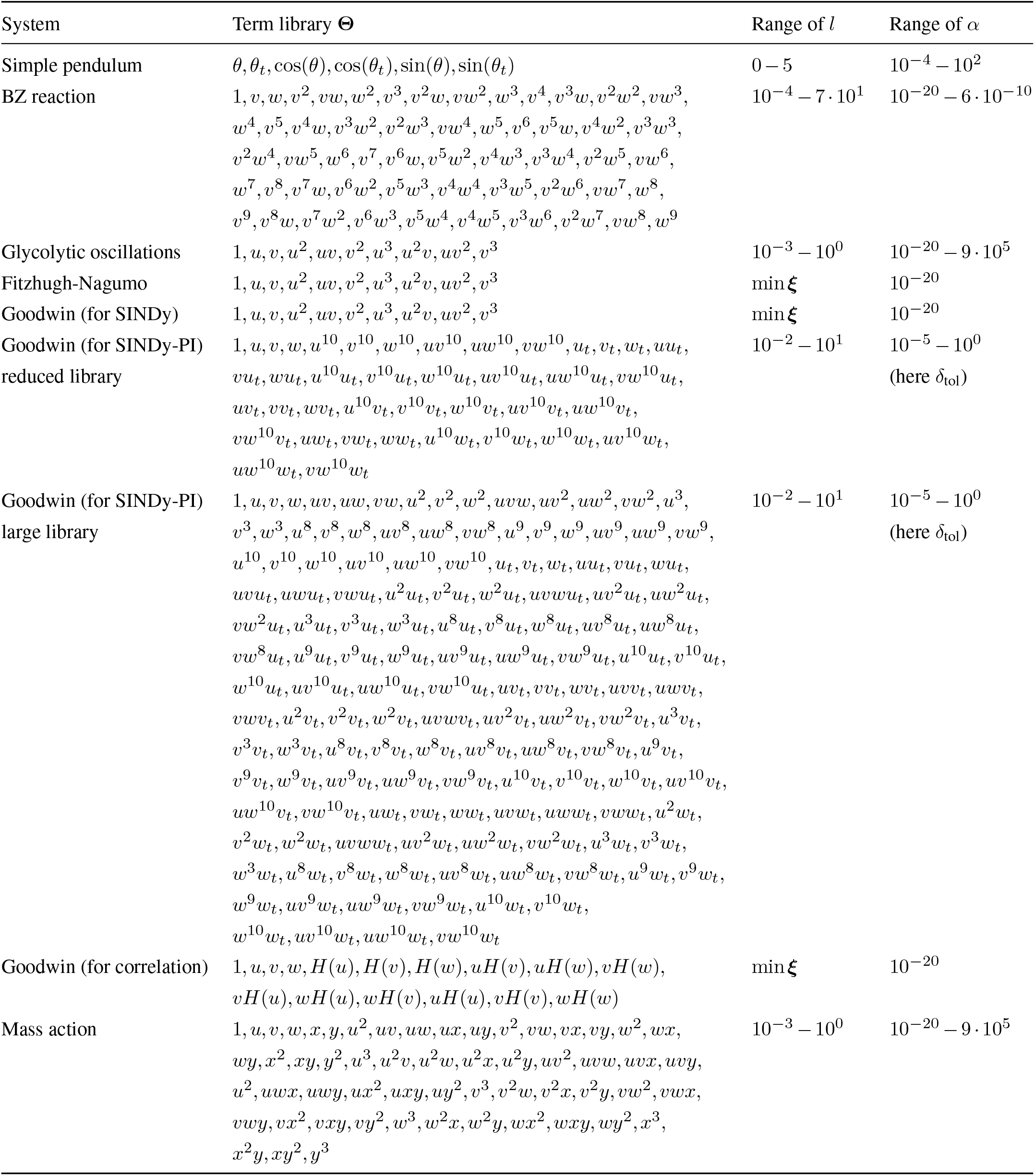
Respective term libraries used for model discovery from experimental data.

#### The SINDy-PI method

The SINDy-PI (parallel implicit) is a further extension of the created implicit SINDy extension. The implicit SINDy extension changes the target of the optimization approach in order to account for rational terms in the model identification (47). An example for Michelis-Menten kinetics is also presented in the examples of the PYSINDY package:

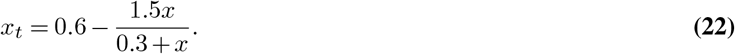

This equation is not solvable for SINDy, but can be translated into an implicit form that then can be solved with SINDy:

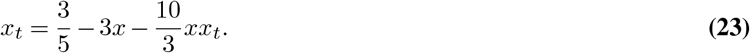

As this formulation is sensitive to noise (47, 96), the parallel implicit extension solves the SINDy regression problem for all terms as the target of the optimization, e.g. *x*_*t*_, *x, xx*_*t*_, ….

#### Libraries used for SINDy

For the analysis of synthetic and experimental data with SINDy, we have used the following combinations of term libraries and hyperparameters listed in Table 3.

#### Evaluation of experimental data

For the evaluation, we look at two parameters: the complexity of the equation and the R^2^ score, which we use as a goodness-of-fit parameter. The R^2^ score is a commonly applied indicator on the goodness-of-fit of a model derived with regression methods. We apply the R^2^ score in the following form,

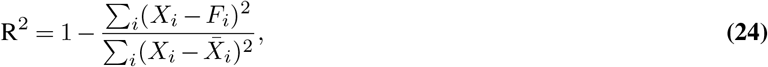

where the numerator expression describes the residual sum of squares and the denominator expression the total sum of squares which are calculated with the original data set *X*_*i*_ and the respective model *X*_*i,t*_ = *F*_*i*_(***X***).

#### Evaluation of synthetic data

We evaluate the performance of the respective SINDy methods by comparing the structure of the candidate models (CM) to the ground true (GT) models provided. We evaluate them under two aspects: if the model has been correctly identified where we accept and relative error in the coefficients ***ξ*** which is less then 2%:

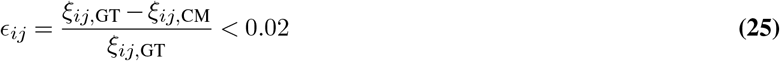

Or if the coefficients lie beyond this 2% threshold but the model still has the same mechanistic form as the ground true model. We evaluate these aspects for model specific sampling ranges where we add Gaussian noise between 1 and 10%.

## Acknowledgements

L.G. acknowledges funding by the KU Leuven Research Fund (grant number C14/18/084 and C14/23/130) and the Research-Foundation Flanders (FWO, grant number G074321N). We also would like to thank Martina Boiardi, Daniel Cebrian-Lacasa and especially Nikita Frolov for useful discussions and comments.

## Author contributions

Conceptualization, B.P. and L.G.; Software, B.P.; Validation, B.P; Formal Analysis, B.P.; Investigation, B.P.; Resources, L.G.; Visualization, B.P; Writing - Original Draft, B.P. and L.G.; Writing - Review & Editing, B.P. and L.G.;Supervision, L.G.; Funding Acquisition, L.G.

## Decleration of Interest

The authors declare no competing interests.

## Footnotes

1 In Halley et al.(87) the authors describe measurement noise in ecology and other biological processes to be pink or 1/f noise which has a linear frequency spectrum on the logarithmic scale, however both noise colors can be distributed normally and the choice of noise color should not impact model identification.

2 Temporal resolution retrieved from personal communication with the authors.

