## Supplementary Information for "From biological data to oscillator models using SINDy"

### Supplementary Note 1: Analysis of reproducibility using SINDy with random seeds

We analyze the how the outcome of analysis is influence by the choice of random seed in the PYTHON script. Choosing a specific random seed can result in randomness of results, however we see when varying the random seed of calculations that the obtained results (here for FHN with $\varepsilon = 0.3, 0.1, 0.01$) still hold (see Fig. S1). This is shown by the distributions which shows minimal amount of points per period required for successful identification for all 50 seeds.

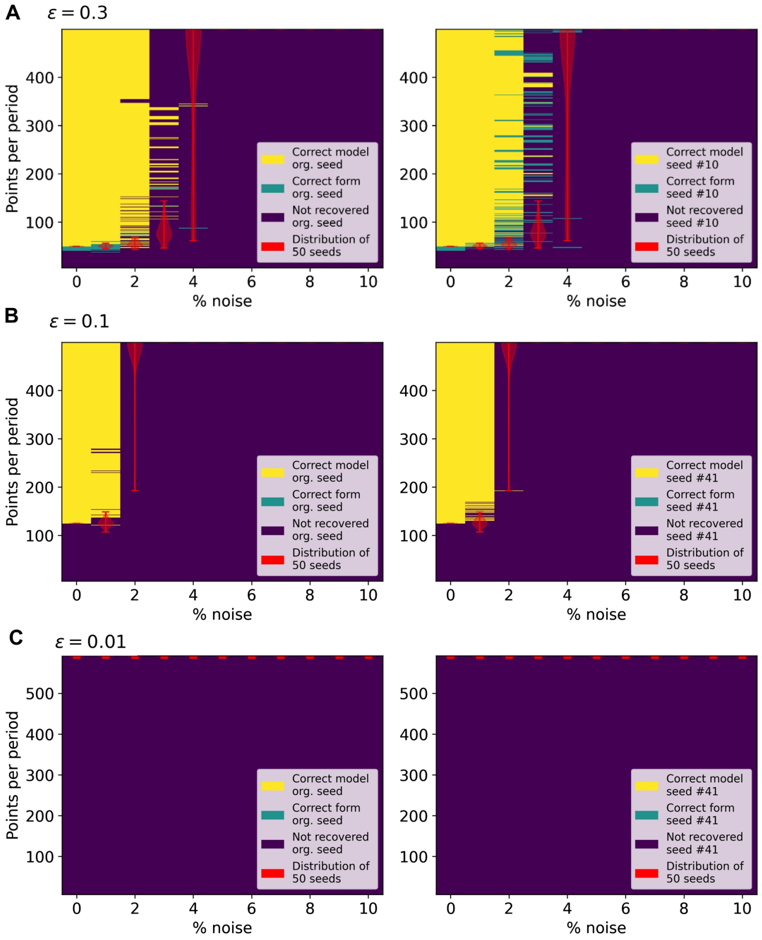


**Fig. S1. Analysis of different random seeds, related to Fig. 3** Setting a single random seed can lead to randomness in results, however running SINDy with 50 different seeds on data from the FHN shows that our obtained results are repeatable independently of the random seed (when compared with results shown in Fig. 3). Red distribution shows the minimal amount of points to achieve correct identification for all seeds as a distribution.**A** Results for FHN with $\varepsilon = 0.3$. **B** Results for FHN with $\varepsilon = 0.1$. **C** Results for FHN with $\varepsilon = 0.01$.

### Supplementary Note 2: Analysis of simple pendulum

We begin with the oscillations of a simple pendulum system from Fig. 1. For this investigation we have generated a video of a swinging brass ball ($m = 44$g) attached to a metal wire ($L = 0.43$m). During the video the ball is observed over a time $t = 12s$ with $dt = 0.008s$ and oscillates 9 times during the observation time. From literature, the equation describing the oscillations of friction-free simple pendulum defined in polar coordinates of the displacement angle is the following:

$$\theta_{tt}(t) = -\frac{g}{L}\sin(\theta(t)) \quad (1)$$

Using this prior knowledge on the dimensions of the system and a simple convolution neural network, which was trained to detect the ball, we track the displacement of the ball over time and calculate the displacement angle $\theta(t)$. Using the original

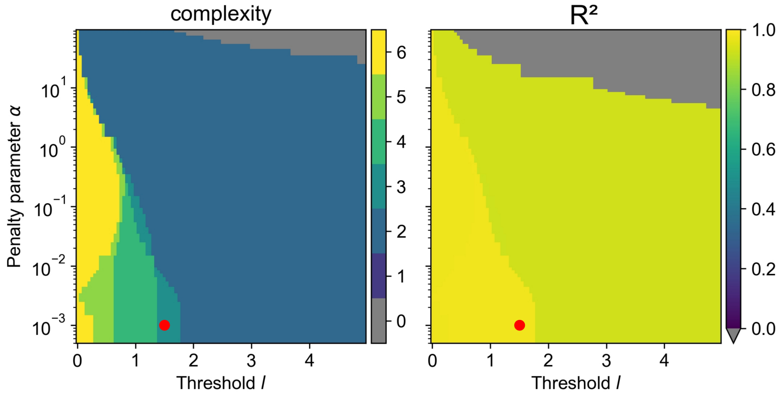


**Fig. S2. Analysis of simple pendulum data, related to Fig. 1** The SINDy algorithm provides a family of models with different complexities depending on the chosen threshold $l$ and penalty factor $\alpha$. Comparing the determined complexity and the $R^2$ score, we select a model resulting at $l = 1.5, \alpha = 10^{-3}$(in red) that is shown in Eq.2.

temporal resolution, we apply the SINDy approach using Ridge regression, scanning over a set of thresholds $l$ and penalizing factors $\alpha$. The results of the analysis can be seen in Fig. S2.
After comparing the determined complexity and $R^2$ score, we select a combination $l = 1.5, \alpha = 10^{-3}$ which combines lowest complexity with the highest adjusted $R^2$ score (marked in Fig. S2). The discovered equation with SINDy and additional application of the small angle approximation leads to the following equations,

$$\begin{aligned} &\theta_{tt} = -24.798\,\theta + 2.071\sin(\theta) - 0.067\sin(\theta_t) \\ &\xrightarrow{\sin(x)\approx x} \theta_{tt} = -22.727\sin(\theta) - 0.067\,\theta_t \end{aligned} \quad \textbf{(2)}$$

This equation has the same form as suggested by theory in Eq. 1 with the additionally discovered friction term, $-b/m\,\theta_t$ with $b$ the damping and $m$ the mass of the ball. Interpreting the parameters as in Eq. 1 and the additional friction term leads to the following:

$$-\frac{g}{L} = -22.727 \rightarrow L = 0.4316\text{m}, \quad -\frac{b}{m} = -0.067. \quad \text{with} \quad m = 44g \rightarrow b = -2.95\frac{g}{[\text{rad}]} \quad \textbf{(3)}$$

We have successfully identified the correct equation from experimental data (with which we are able to determine the correct length of the pendulums wire) and also discovered the realistic friction term using the SINDy algorithm. Therefore, as a proof-of-concept in the case of high-resolution, low-noise and prior-knowledge of the mechanism, we are able to identify the underlying governing equations correctly.

### Supplementary Note 3: Application on BZ reaction data

**A. Data correction of original potentiometric data.** In order to hopefully rediscover the underlying Oregonator model from BZ data, we detrended the data using a rolling window mean and additionally rescaled all local maxima and minima of the periods to the maximum and minimum of the first period. With this we wanted to reduce the impact of reagent depletion in the reaction, which causes the levels of bromine ions and cerium to decrease over time. Furthermore to enable the identification of the Oregonator model, we used prior knowledge on the model and adjusted the data according to the original position of the limit cycle from the Oregonator model.

**B. Analysis.** Now we can consider the well-studied chemical oscillatory system, the Belousov–Zhabotinsky reaction, one of the earliest studied chemical systems is the Bromate-Cerium-Malonic Acid system, which also led to the formulation of a mechanism describing the emergence of temporal oscillations, called the Field-Kórós-Noyes (FKN) mechanism (1). In this system the main interaction occurs between three compounds: the negatively charged bromine compounds or bromine ions $[Br^-]$, the positively charged cerium(IV) and cerium(III) ratio $[Ce^{4+}]/[Ce^{3+}]$ and the neutral bromous acid $[HBrO_2]$ (1). Together with potentiometric measurements of bromine and cerium ion concentrations, FKN determined a reduced mechanism that was able to describe the emergence of oscillations. Later, Fields and Noyes have shown that the FKN mechanism can be reduced even further to a set of three ordinary differential equations, also called the Oregonator (2) which brought into the following form by Tyson (10, 11):

$$\begin{aligned} u_t &= p\left[u(1-u) - v(u-q)\right], \\ v_t &= p'(uv + gv - 2fw), \\ w_t &= u - w. \end{aligned} \tag{4}$$

Here, $u, v, w$ can be assigned to the mentioned compounds, being $u = [\text{HBrO}_2]$, $v = \left[\text{Br}^-\right]$ and $w = \left[\text{Ce}^{4+}\right] / \left[\text{Ce}^{3+}\right]$. However, the concentration of component $[\text{HBrO}_2]$ is not obtainable, as it can not be measured potentiometrically.
Therefore, we investigate the reduced system, which means to simplify the Oregonator model in Eq. 4 with a steady-state approximation of the $u_t$ equation. Reducing the Oregonator model is commonly done when investigating its dynamical behavior, see e.g (9, 10, 12) in which different equations of the system are approximated as a steady state. Now reducing the system in Eq. 4 with $u_t = 0$, leads to the following reduced system,

$$\begin{aligned} v_t &= p'(uv + gv - 2fw), \\ w_t &= u - w. \\ \text{with } u(v) &= \frac{(1-v) + \sqrt{(v-1)^2 - 4q}}{2}. \end{aligned} \tag{5}$$

As we see the expression of $u(v)$ is not trivial, but what does this mean for the application of SINDy? Assuming, that we do not have this knowledge on the underlying mechanism, applying SINDy naively with a term library containing combinations of $v$ and $w$ up to the fifth order is able to produce models which tend to blow-up. We identify the equations by checking all possible solutions in for different pairs of threshold $l$ and penalty factor $\alpha$ for polynomial libraries of order up to 6 (see Fig. S3). We adjusted the parameters of the SINDy approach carefully, aiming on providing models which do not result in a blow-up and we were only able to identify such a model at $l = 0.04, \alpha = 10^{-20}$ with a library containing combinations up to the sixth order:

$$\begin{aligned} v_t &= f(v, v^2, vw, v^3, \ldots, vw^4), \\ w_t &= f(v, w, v^2, vw, w^2, v^3, \ldots, v^2w^3, vw^4, w^5) \end{aligned} \tag{6}$$

When compared to the original time series and the limit cycle of the BZ reaction oscillations in Fig. 1, the model is able to reproduce the behavior. With knowledge of the mechanism and also the reduction of the system, we are able to identify that the complexity of the discovered model expresses a Taylor series expansion of the steady state. Although not easily interpretable, the SINDy algorithm was able to quantitatively reproduce the temporal oscillations of the BZ reaction just from experimental data. The oscillations have a similar amplitude, while the period is different as the discovered model oscillates slower than the experimental data. The most striking difference can be seen when comparing the resulting limit cycle to the experimental limit cycle of cerium and bromine: slow changes of both compounds are captured well by the model. However, when the concentration of one compound changes rapidly compared to the other it leads to sharp corners in the limit cycle. These changes are not captured by the discovered model. This shortcoming can be directly linked to the reduction of the underlying system to only two compounds: As comparing the Taylor expansion shows, we only approximate the rapid changes introduced through the behavior of bromous acid or component $u$. Therefore, it becomes clear that not being able to observe all variables of a experimental system can significantly obstruct the correct identification and disallow for an interpretable model, despite capturing the qualitative behavior of the system.

**C. Inferred reduced model of the BZ reaction.** To improve readability of the main text, we show here the results of the application of SINDy on the original data after transformation.

$$\begin{aligned} v_t =& 0.392v + 37.79 \cdot 10^3 v^2 - 3.682vw - 33.5 \cdot 10^9 v^3 + 1.29 \cdot 10^6 v^2 w - 76.3vw^2 + 23.86 \cdot 10^6 v^2 w^2 \\ & - 129.04vw^3 + 72.37 \cdot 10^6 v^2 w^3 + 447.32vw^4, \\ w_t =& 66.8v + 0.003w + 649.03 \cdot 10^6 v^2 - 262.95 \cdot 10^3 vw + 1.84w^2 - 228.81 \cdot 10^3 v^3 + 9449 \cdot 10^9 v^2 w \\ & - 533.88 \cdot 10^3 vw^2 - 4.63w^3 + 35.57 \cdot 10^6 v^3 w + 50.76 \cdot 10^9 v^2 w^2 + 12.82 \cdot 10^3 vw^3 \\ & - 186.82w^4 - 3.36 \cdot 10^6 v^3 w^2 - 25514. \cdot 10^9 v^2 w^3 + 39.72 \cdot 10^6 vw^4 - 606.31w^5 \end{aligned} \tag{7}$$

The polynomial consists of high order terms that aim on accounting for the reduction of a system by approximating it with a Taylor-series expansion:

$$u(v) \approx \frac{1}{2}(\sqrt{1-4q}+1) + \frac{1}{2}\left(\frac{-1}{\sqrt{1-4q}} - 1\right)v - \frac{qv^2}{(1-4q)^{3/2}} - \frac{qv^3}{(1-4q)^{5/2}} - \frac{q(q+1)v^4}{(1-4q)^{7/2}} + \mathcal{O}(v^5) \tag{8}$$

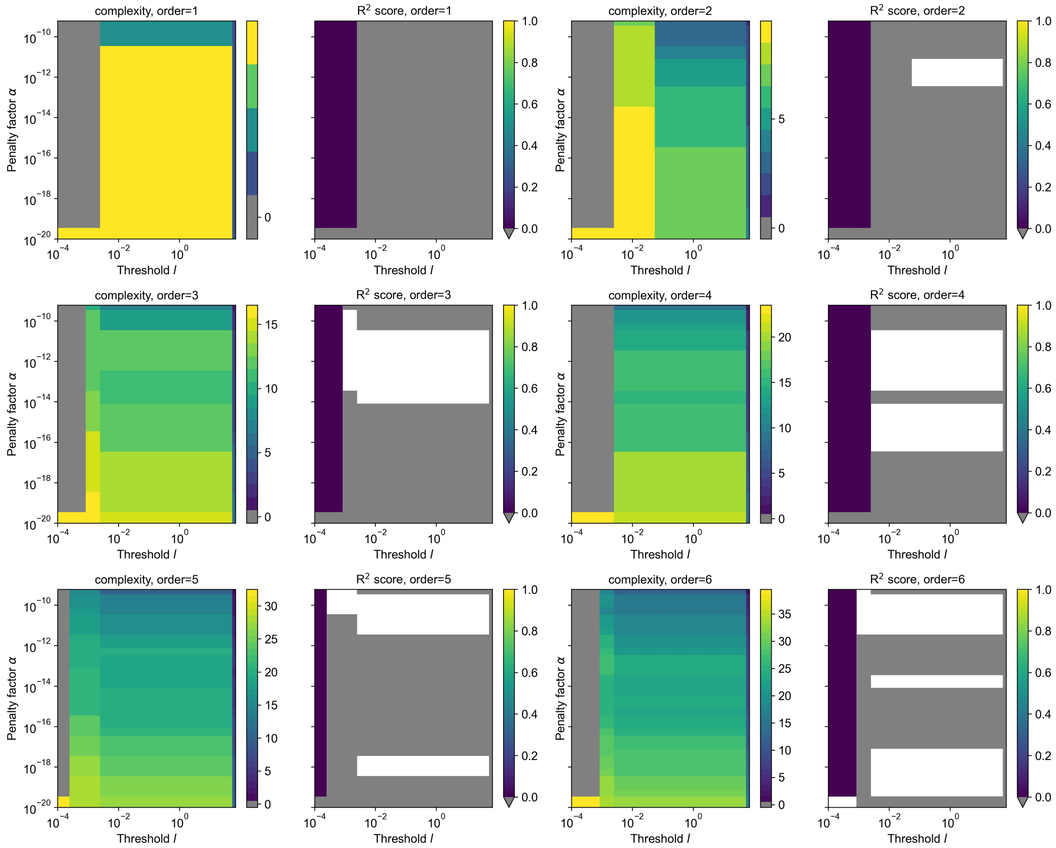


**Fig. S3. Analysis of BZ reaction data, related to Fig. 1** Averaged BZ time series which eliminates sharp change and variability of the limit cycle, analyzed for different sets of libraries with their respective complexity and $R^2$ score. If areas are white the identified models either did not converge or show $R^2$ values of less than $0$.

### Supplementary Note 4: Study of noise filtering in low-data, high-noise situations for model recovery

In order to understand the differences in the success of model identification in Fig. 3D, we investigated the influence of sampling on the form of noise and performance of the studied noise filtering techniques low-pass filter, Wiener filter, LPSA and Savitzky-Golay filter in Fig. S4. The formulations of the noise filtering techniques are taken from Kantz et al.(3) for the low pass filter, Wiener filter and LPSA, and for the Savitzky-Golay filter we reused the code provided in Lejarza et al.(4).

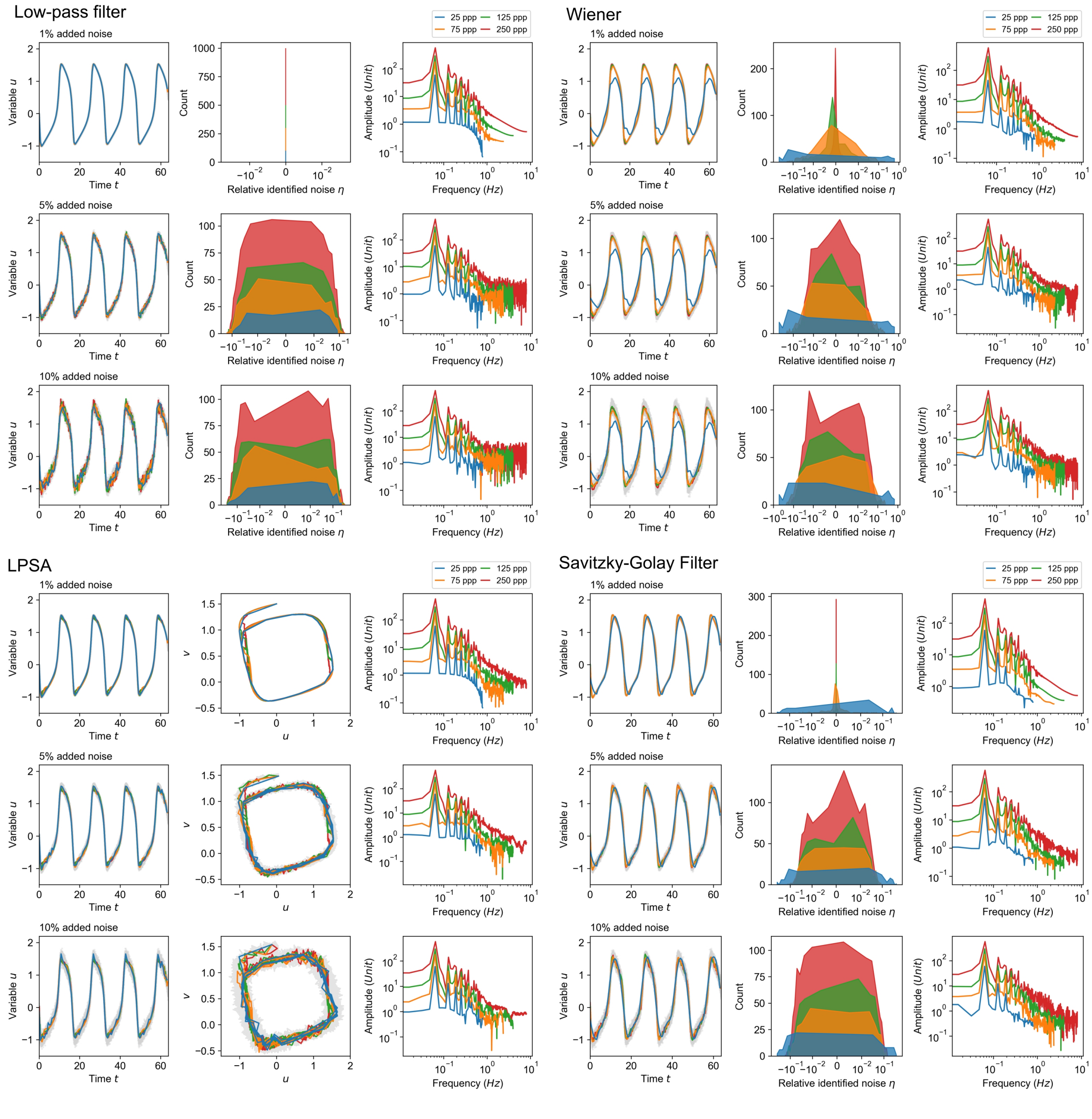


**Fig. S4. Investigation on the role of subsampling and form of noise for different noise filtering techniques, related to Fig. 3** **Low-pass filter:** When low noise levels are introduced different sampling does not have an impact on the signal and any additional noise is filtered completely. However, the higher the noise levels become the more the noise distribution becomes either uniform (for 5%) or degenerates (for 10%), requiring precise adjustment of the cutoff frequency. **Wiener filter:** The Wiener filter is able to extract low-level noise (1%), however for small sampling the required from of noise (normally distributed) degenerates and the filter fails. This is also the case for higher noise levels, where the identified noise is not normally distributed, thus obstructing noise filtering with the Wiener filter. **LPSA:** The local phase space averaging is able to handle low noise situations, however struggles when noise is increased to account for the introduced variability in the signal. **Savitzky-Golay filter:** The low order polynomial approximation in moving windows reduces the filtering performance as the higher the variability, the more precise must be the window size on which the regression is applied. Otherwise, the regression overestimates the variability of the signal, resulting in a locally incorrect representation of the signal.

### Supplementary Note 5: Multiple trajectories and strong time scale separation

We show here the application of the multiple trajectories setup explained in the section 'Addressing noise and low-data in equidistant sampling with multiple trajectories' on data from the FHN with a time scale separation $\varepsilon = 0.1$.
Similarly to the case shown in Fig. 4C we generate data sets with different initial conditions (IC) and try to identify the model form this aggregated data. We find that providing more initial conditions requires high temporal resolution of the data and is only able to identify the correct form of FHN model for noise levels (see Fig. S5). Here the averaging of the solution over multiple trajectories underestimates the role of time scale separation in the model and drops terms that are included in the ground truth solution.

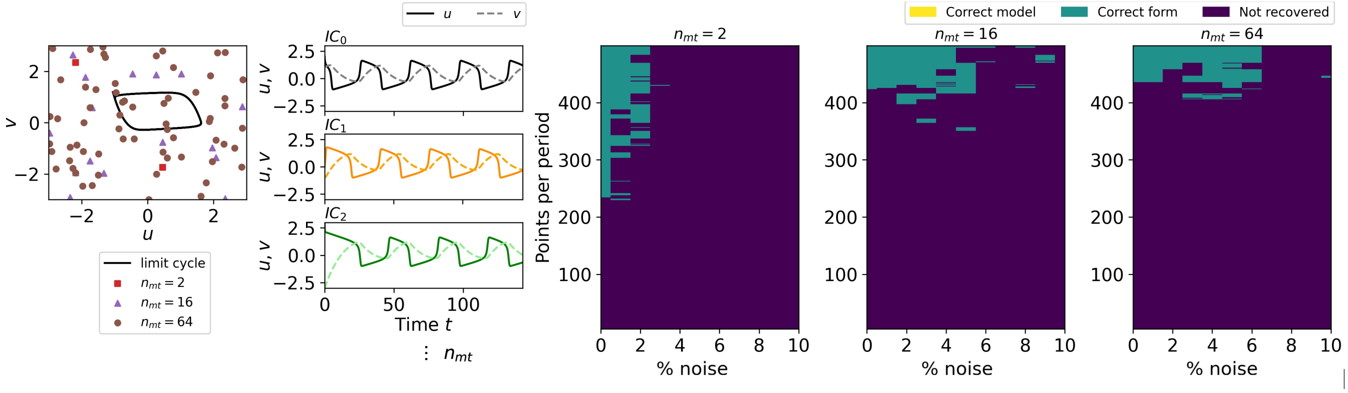


**Fig. S5. Increasing data amounts with multiple trajectories fails for stronger time scale separation $\varepsilon = 0.1$, related to Fig. 4** Providing more trajectories of the underlying dynamics with different, random initial conditions (IC) does not improve identification of the correct form of the FHN equation. Identifying the correct form is only possible for high resolution and low noise, as averaging over multiple trajectories underestimates the impact of time scale separation in the identified model. The different IC are shown in the left panel: squares for two ICs, triangles for 16 ICs and circles for 64 ICs.

### Supplementary Note 6: Non-polynomial nonlinearity in model formulation - Hill function

We investigate the performance of SINDy for other forms of nonlinearity in the model formulation with the Hill-function. In our analysis, we provide a term library containing terms up to the third order and try to approximate the dynamical behavior of the Goodwin model with $n = 10$. Results of the SINDy approximation for $n = 8$:
Results of the SINDy approximation for $n = 10$:

$$\begin{aligned}
u_t =& 4.551u + 52.994u^2 + 2.016uv - 3.663uw - 0.104v^2 + 69.809u^3 - 66.216u^2v \\
& - 42.768u^2w + 37.525uv^2 - 5.023uvw + 0.894uw^2 + 0.657v^3, \\
v_t =& -3.816u + 32.646u^2 + 12.819uv + 5.468uw - 2.519v^2 - 36.703u^3 \\
& - 39.648u^2v - 24.347u^2w + 6.167uv^2 - 3.945uvw - 1.718uw^2 + 2.115v^3 + 0.761v^2w, \\
w_t =& 6.798u + 0.796v - 0.105w - 178.464u^2 + 45.158uv - 9.437uw - 1.516v^2 \\
& + 0.380vw + 881.399u^3 - 429.518u^2v + 117.386u^2w + 52.729uv^2 \\
& - 29.157uvw + 3.235uw^2 + 0.580v^2w - 0.110vw^2,
\end{aligned} \tag{9}$$

The resulting model usually contains a large fraction of the suggest terms in the polynomial library, thus making them challenging to interpret yet are able to approximate the behavior of the system well (see Fig. 6A).
When SINDy-PI is applied, we are able to identify the nonlinear approximation of the Hill-function, $1/w^{10}$ (see Fig. S6A), when provided data has sufficient resolution and low-noise, and we provide a term library that only provides relevant terms already included in the underlying model (see Fig. 6B,C). We also identify equations for the $w_t$ equation shown in Table 2 which despite their more complicated form are approximations of the dynamics (see Fig. S6B). We also tested SINDy-PI with a larger library, for which we are not sure about the structure of the model, still using high resolution (398 ppp) and no noise conditions. Here, similarly to original SINDy the identification suffers from selecting too many possible terms, however it is able to provide functions for all chosen hyperparameter values (see Fig. S7). However, all the provided models fail when evaluated numerically which shows that even when using SINDy-PI to provide nonlinear terms for the original SINDy, not only resolution and noise mitigation play an important role but also the correct truncation of the library.

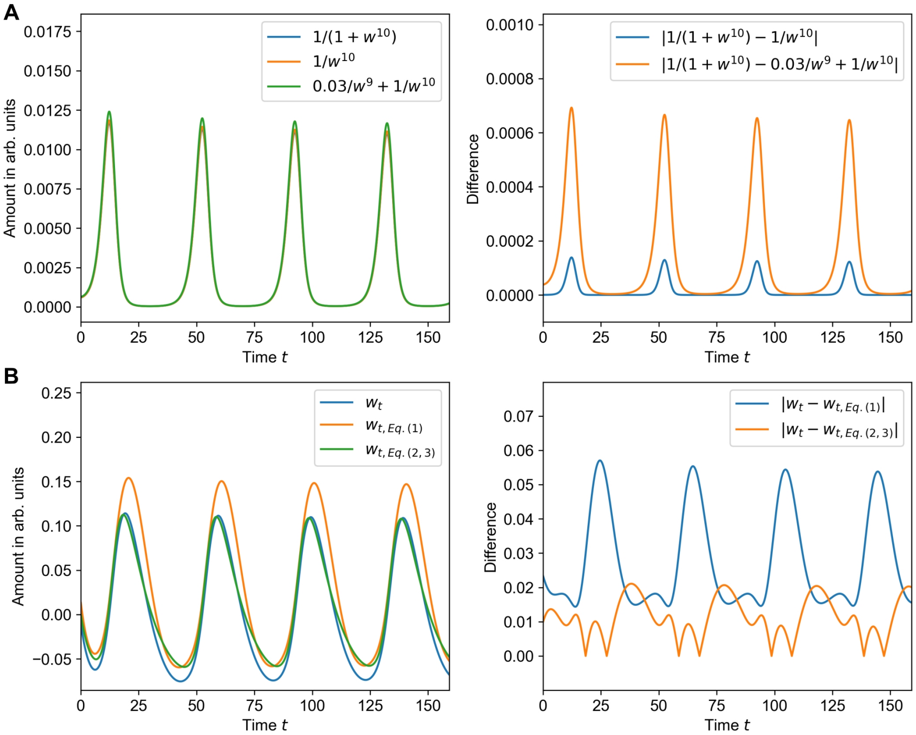


**Fig. S6. SINDy-PI is able to identify approximations of the Hill-function and $w_t$ equation from data, related to Fig. 6 A** SINDy-PI is able to identify two close approximations of the Hill-function in the equation $u_t$. **B** For the equation $w_t$, SINDy-PI does not identify the simplest representation, but is able to identify a more complex approximation of the dynamical behavior.

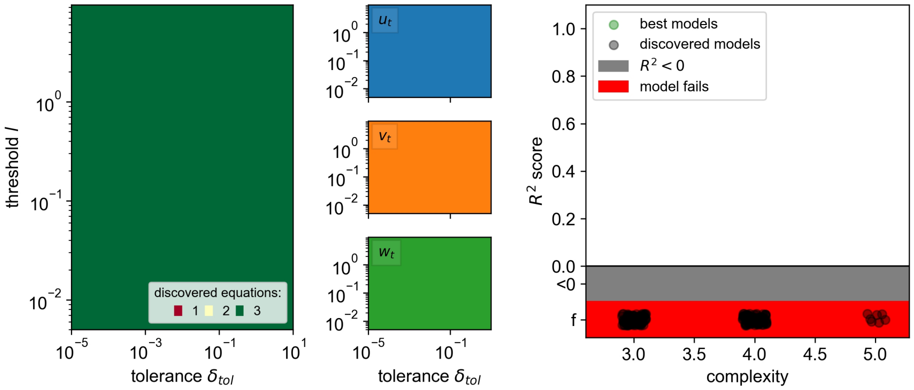


**Fig. S7. Increasing amount of terms in SINDy-PI library provides failing models, related to Fig. 6** When more terms are provided to the term library of SINDy-PI and the algorithm is applied on data with high resolution (398 ppp) and no noise, SINDy-PI is able to identify the full set of equations for all chosen hyperparameters. However, when analysing the equations numerically using the $R^2$ score, all identified models fail to be simulated and are not able to reproduce the provided data.

### Supplementary Note 7: Analysis of different dimensions in mass action model

In section 'High dimensionality in the mass action oscillator', we investigate how dimensionality reduction can enhance model identification with SINDy and here we provide the models identified with the use of Fig. 7C and are mentioned in the section. For $(u,v)$ and $l = 10^{-2}, \alpha = 10^{-2}$ we get the following equation:

$$\begin{aligned} u_t &= 0.377u + 0.061u^2 - 6.902u^3 + 0.277u^2 v, \\ v_t &= -3.414u - 0.044v - 0.094uv. \end{aligned} \quad (10)$$

For the other equations $(v,w)$ at $l = 10^{-2}, \alpha = 10^{-2}$ we identify that:

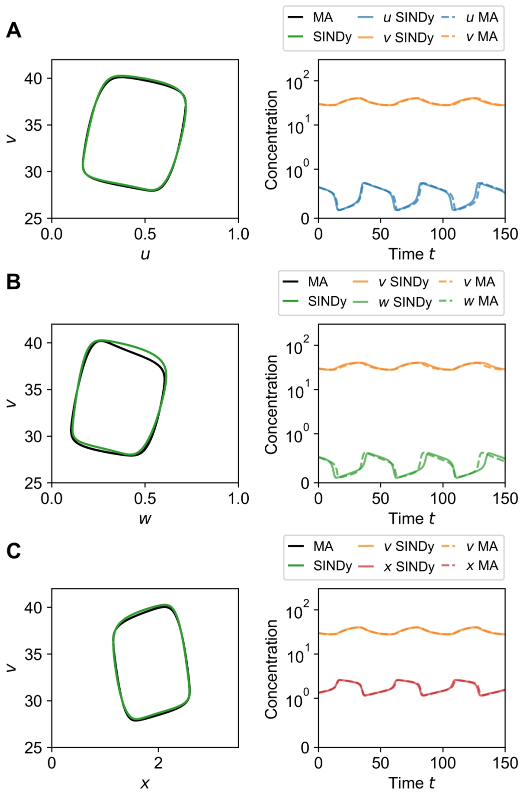


**Fig. S8. Evaluation of inferred models from dimension reduction, related to Fig. 7** Simulated time series of the models Eq. 10 in **A**, Eq. 11 in **B** and Eq. 12 in **C**,

$$
\begin{aligned}
v_t &= -0.1491 - 0.055v - 4.107w - 0.129vw \\
&\quad + 2.347w^2 + 0.025vw^2 + 7.377w^3, \\
w_t &= 0.146vw^2 - 0.193w^3,
\end{aligned} \tag{11}
$$

and for $(v, x)$ at $l = 2 \cdot 10^{-2}, \alpha = 10^{-19}$ that:

$$
\begin{aligned}
v_t &= -0.0661 - 0.065v + 1.578x + 0.048vx \\
&\quad + 0.125x^2 - 0.650x^3, \\
x_t &= -0.036v + 0.711x - 1.654x^3,
\end{aligned} \tag{12}
$$

are able to approximate the corresponding behavior in the reduced system (see Fig. S8B and C).

### Supplementary Note 8: Analysis of glycolytic oscillations following step-by-step instructions

The data of NADH and ATP has been extracted from Özalp et al.(7), Fig. 2. The data has a resolution of $dt = 1s$ according to personal communication with the authors, and is adjusted for drift, however the time series data is already cleaned from noise in Özalp et al.(7) but reused in Fig. 13.2a in Olsen et al.(6) with noise. From Fig. 13.2a in Olsen et al.(6) we evaluated that the noise levels present are around 20% of the normal distribution of the data and we therefore add this amount of noise to the experimental data following Eq. 11. We take only a part of the full data set and apply SINDy on it, shown in Fig. S9A before and B after noise filtering. Afterwards, we center and normalize the data set and apply SINDy again, with (Fig. S9C) and without noise (Fig. S9D), which lead to the derivation of Eq. 16.
For the case of high resolution, we have taken the data set shown in Özalp et al.(7) in Fig.2 and sampled the data with higher resolution of $dt_1 = 0.25$ (380 ppp, ten times higher resolution), which we examine after detrending and rescaling (step (1)). In order to resemble experimental data we add 20% of Gaussian noise with the same characteristics as the lower resolution data. We then go through the same steps while showing the result at every step: high resolution ($dt = 0.25s$, Fig. S9E with noise,

Fig. S9F without noise) and very high resolution ($dt = 0.01$, Fig. S9G with noise and Fig. S9H without noise) which lead to Eqs. 13 and 17.
With this increase, we were able to identify the following model at $\alpha = 1 \cdot 10^{-17}$ and $l = 1 \cdot 10^{-2}$ with $R^2 = 0.7824$ and complexity $k = 12$:

$$\begin{aligned} u_t =& -0.023 - 0.018u - 0.274v + 0.032v^2 \\ & -0.081u^3 + 0.124v^3, \\ v_t =& 0.019 + 0.188u + 0.076v + 0.030uv \\ & -0.014v^2 - 0.027u^3 \end{aligned} \tag{13}$$

This model with a reduced complexity preserves the previously identified dynamical behavior, comparable to the one with very high resolution in Fig. 10 (data not shown here).

### Supplementary Note 9: Initial study on performance of Weak-SINDy for oscillatory systems

We have also analyzed in an initial study how the Weak-SINDy method (5, 8) performs when applied on data from the FHN model with different levels of time scale separation (see Fig. S10). As the the Weak SINDy does not use any derivatives but the integral form of the SINDy optimization problem, it also shows improved performance for strong time scale separation.

### Bibliography SI

1. Richard J. Field, Endre Koros, and Richard M. Noyes. Oscillations in Chemical Systems. II. Thorough Analysis of Temporal Oscillation in the Bromate–Cerium–Malonic Acid System. *Journal of the American Chemical Society*, 94(25):8649–8664, 1972.
2. Richard J. Field and Richard M. Noyes. Oscillations in chemical systems. IV. Limit cycle behavior in a model of a real chemical reaction. *The Journal of Chemical Physics*, 60(5):1877–1884, mar 1974.
3. Holger Kantz and Thomas Schreiber. *Nonlinear time series analysis*, volume 7. Cambridge university press, 2004.
4. Fernando Lejarza and Michael Baldea. Data-driven discovery of the governing equations of dynamical systems via moving horizon optimization. *Scientific Reports*, 12(1):1–15, jul 2022.
5. Daniel A. Messenger and David M. Bortz. Weak SINDy: Galerkin-Based Data-Driven Model Selection. *Multiscale Modeling & Simulation*, 19(3):1474–1497, jan 2021.
6. Lars Folke Olsen and Anita Lunding. Oscillations in Yeast Glycolysis. In *Understanding Complex Systems*, pages 211–224. Springer Science and Business Media Deutschland GmbH, 2021.
7. Veli C. Özalp, Tina R. Pedersen, Lise J. Nielsen, and Lars F. Olsen. Time-resolved Measurements of Intracellular ATP in the Yeast Saccharomyces cerevisiae using a New Type of Nanobiosensor. *Journal of Biological Chemistry*, 285(48):37579–37588, nov 2010.
8. Patrick A. K. Reinbold, Daniel R. Gurevich, and Roman O. Grigoriev. Using noisy or incomplete data to discover models of spatiotemporal dynamics. *Physical Review E*, 101(1):010203, jan 2020.
9. John J. Tyson. Analytic representation of oscillations, excitability, and traveling waves in a realistic model of the Belousov–Zhabotinskii reaction. *The Journal of Chemical Physics*, 66(3):905–915, feb 1977.
10. John J. Tyson. Oscillations, Bistability and Echo waves in models if the Belousov-Zhabotinskii reaction. *Annals of the New York Academy of Sciences*, 316(1):279–295, feb 1979.
11. John J. Tyson. From the Belousov–Zhabotinsky reaction to biochemical clocks, traveling waves and cell cycle regulation. *Biochemical Journal*, 479(2):185–206, jan 2022.
12. John J. Tyson and Paul C. Fife. Target patterns in a realistic model of the Belousov-Zhabotinskii reaction. *The Journal of Chemical Physics*, 73(5):2224–2237, 1980.

$LN@col2$

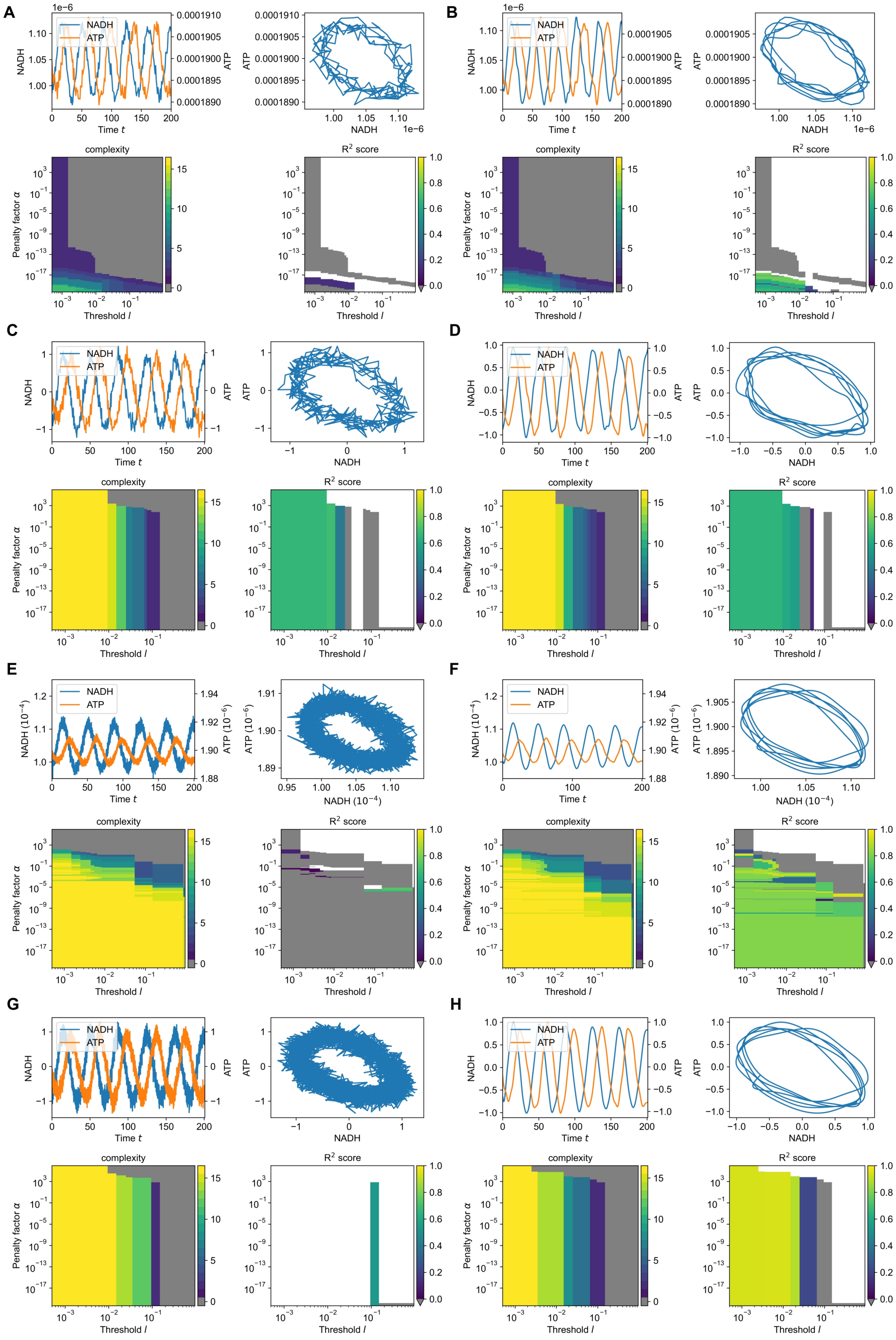


**Fig. S9. Analysis of the glycolysis data set from Özalp et al.(7) with interpolation, related to Figs. 9,10 A and B** Naive application of SINDy on the original data without and with noise filtering, does not lead to interpretable models. **C** Rescaling and normalizing the data allows to identify models with high complexity. **D** Following our investigation of noise filtering, we apply a suitable filtering method and are able to derive a model with complexity $k = 15$ as shown in Eq. 16. **E** In order to improve analysis, we interpolate the data between the measured data points and reduce complexity of the underlying models. **F** After noise filtering, we select a model with the best trade off between $R^2$ score and complexity $k$ shown in Eq. 13. **G** Increasing the data even more allows to reduce the complexity and increase the $R^2$ score. **H** After additional noise filtering, we are able to achieve an interpretable and analyzable model in Eq. 17.

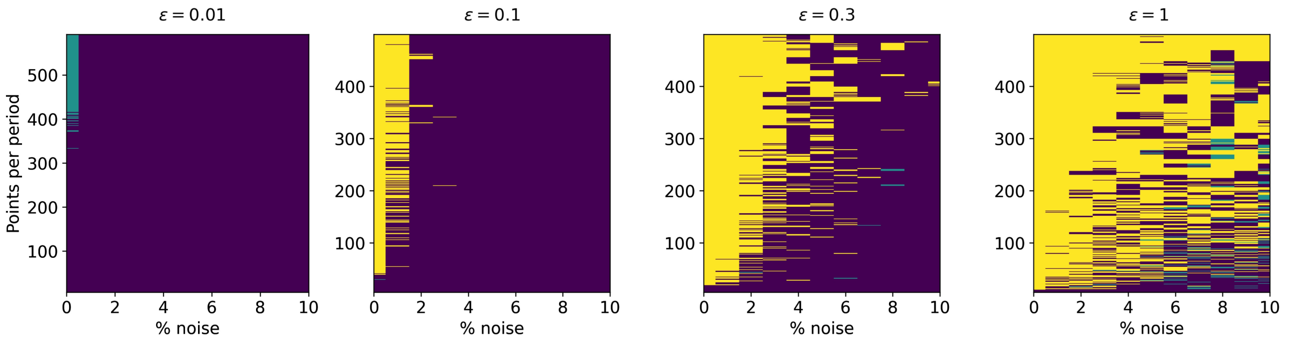


**Fig. S10. Analysis of Weak SINDy performance for different time scale separation in the FHN model, related to Fig. 3** The analysis shows improved identification for even strong time scale separation in the FHN model compared to the original, derivative SINDy method. Applying Weak SINDy poses an intersting further line of work.
